# Understanding the interplay of sleep and aging: Methodological challenges

**DOI:** 10.1101/713552

**Authors:** Beate E. Muehlroth, Markus Werkle-Bergner

## Abstract

In quest of new avenues to explain, predict, and treat pathophysiological conditions during aging, research on sleep and aging has flourished. Despite the great scientific potential to pinpoint mechanistic pathways between sleep, aging, and pathology, only little attention has been paid to the suitability of analytic procedures applied to study these interrelations. On the basis of electrophysiological sleep and structural brain data of healthy younger and older adults, we identify, illustrate, and resolve methodological core challenges in the study of sleep and aging. We demonstrate potential biases in common analytic approaches when applied to older populations. We argue that uncovering age-dependent alterations in the physiology of sleep requires the development of adjusted and individualized analytic procedures that filter out age-independent inter-individual differences. Age-adapted methodological approaches are thus required to foster the development of valid and reliable biomarkers of age-associated cognitive pathologies.

## Introduction

In search of novel biomarkers and treatment targets for age-related pathologies, sleep has recently attracted much attention. Disrupted sleep represents one of the earliest symptoms of Alzheimer’s disease (Lim et al., 2013; Lucey et al., 2019), but might also potentiate, accelerate, or even cause cognitive pathology in old age (Shokri-Kojori et al., 2018; Vaou et al., 2018). Because of the assumed bidirectional relationship between sleep and Alzheimer’s pathology (Ju et al., 2014; Mander et al., 2016; Noble and Spires-Jones, 2019; Vaou et al., 2018; Winer et al., 2019), studying the interrelations between sleep and neurocognitive aging has become increasingly popular.

Even in the absence of any diagnostically identifiable pathology, healthy aging entails fundamental yet very characteristic changes in sleep (Mander, 2017; Ohayon et al., 2004; Vitiello, 2006). Overall, older adults’ sleep is shifted earlier in time (including earlier bed and wake times), gets lighter, more fragile and shorter (Dijk et al., 2000; Monk et al., 2005). Sleep alterations are evident in the global distribution and succession of sleep stages (i.e., the *macro-structure* of sleep; Feinberg, 1974; Ohayon et al., 2004), but also involve a changed signature of electrical oscillations that define and characterize different sleep states (i.e., the *micro-structure* of sleep; cf. Box 1; Carrier et al., 2011; Crowley et al., 2002). In particular, older adults’ sleep is characterized by a drastic reduction of deep non-rapid eye movement (NREM) sleep, so-called slow-wave sleep (SWS), that is accompanied by an increase of light NREM sleep (Carrier et al., 2011; Danker-Hopfe et al., 2005; Ohayon et al., 2004; Redline et al., 2004). In contrast to NREM sleep, age-related changes in the amount of rapid eye movement (REM) sleep are typically more subtle (Redline et al., 2004; Scullin and Gao, 2018), although a pronounced reduction of rapid eye movements within this sleep stage has been reported (Darchia et al. 2003). Moreover, with advancing age, high-amplitude slow oscillations (< 1 Hz), slow delta waves (1–4 Hz), and discrete sleep spindles (11–16 Hz) defining NREM sleep appear less often and with reduced amplitudes and altered topography (Crowley et al., 2002; Dubé et al., 2015; Fogel et al., 2012; Landolt and Borbély, 2001; Landolt et al., 1996; Martin et al., 2013). Diminished homeostatic sleep pressure, circadian shifts, dysregulation of neurotransmitters, and structural brain alterations may contribute to these pronounced changes in sleep during aging (for more details see Dijk et al., 2000; Mander et al., 2017; Monk et al., 2005; Skeldon et al., 2016; Zhong et al., 2019).

Standard criteria to describe these age-related alterations in sleep physiology are only rarely tested and validated within heterogeneous samples that include older adults (but see, e.g., Ujma et al., 2015, 2019; Warby et al., 2014). Therefore, in the following, we aim to demonstrate the possible benefits and limitations of applying established indicators of sleep physiology in the context of aging research. We aim to raise awareness for the complexity of aging that entails widespread structural and functional brain changes that affect sleep in multiple ways (Dubé et al., 2015; Fogel et al., 2012, 2017; Muehlroth et al., 2019b). To identify meaningful biomarkers that can explain or even predict cognitive decline in old age, we need to promote the use of sensitive and age-adjusted methodology that captures within-person age-dependent changes in the physiology of sleep, rather than age-inflated inter-individual differences (Lindenberger et al., 2011).

### Preamble: How do we define ‘old age’ and ‘aging’?

As an inevitable prerequisite, the identification of changes in sleep physiology during aging requires an appropriate and transparent rationale on how research should delineate and investigate ‘old age’ and ‘aging.’ The literature in (cognitive) neuroscience is dominated by cross-sectional group comparisons of ‘younger’ (∼20–30 years) and ‘older’ adults (∼60–80 years) (Browning and Spilich, 1981; Hedden and Gabrieli, 2004). This is grounded in the notion that the brain’s structure and function has already undergone profound alterations when humans reach their 60’s (e.g., Li et al., 2004; Ohayon et al., 2004; Raz et al., 2005; Rönnlund et al., 2005; Ziegler et al., 2012). Beyond stressing the importance of a general agreement on the definition of ‘old age’ to ensure the comparability of research on aging, we want to highlight two challenges that accompany such a definition: First, the *chronological* age of individuals can deviate from their *biological* age. Second, age *differences* derived from such group comparisons cannot capture aging in terms of age-related *change*.

First and foremost, one should bear in mind that *chronological* age, i.e., the time that has passed since birth, is a variable that does not induce or explain any developmental change in itself (Wohlwill, 1970). At best, it can serve as a proxy for an individual’s expected functional capacity that relies on true mechanistic alterations taking place as humans age (Li and Schmiedek, 2002; MacDonald et al., 2011). The conceptual and methodological pitfalls of using chronological age as an index of development or aging have been discussed elsewhere in detail (e.g., Cole et al., 2017a, 2019; Lindenberger and Pötter, 1998; MacDonald et al., 2011; Wohlwill, 1970). Due to great inter-individual variation in the manifestation of aging effects (Cabeza et al., 2018; Habib and Nyberg, 2007; Lindenberger, 2014), particularly in old age, chronological age can deviate greatly from an individual’s biological age (Cole et al., 2017a, 2019; MacDonald et al., 2011; Steffener et al., 2016). Importantly, physiological brain age, as for instance derived from estimates of structural brain integrity, could represent a better and more informative predictor of differences in aging, individual (brain) health, functional capacity, and mortality (Burzynska et al., 2013; Cole et al., 2018, 2017a, 2017b; Franke et al., 2010). Notably, also brain age derived from the sleep electroencephalogram (EEG) has lately shown its potential to serve as a sensitive index of aging and pathology (Sun et al., 2019). In the course of this paper, we will suggest that the sensitivity of sleep physiology as a potential biomarker of brain age, aging, and age-related disease is closely tied to a fair and adapted investigation and evaluation of sleep in aged individuals.

Strictly speaking, the term aging describes a *process of change*, taking place as organisms grow older. Cross-sectional age comparisons are not appropriate to inform us about these changes (Hertzog and Nesselroade, 2003; Hofer and Sliwinski, 2001; Li and Schmiedek, 2002; Lindenberger and Pötter, 1998; Lindenberger et al., 2011; Overton, 2010; Raz and Lindenberger, 2011). When sampling age-heterogeneous groups cross-sectionally, essential confounds such as cohort effects or sampling bias may covary with age and thus prevent or bias conclusions on intra-individual dynamics of change (Hertzog and Nesselroade, 2003; Hofer and Sliwinski, 2001; Li and Schmiedek, 2002; Lindenberger et al., 2011; Rönnlund et al., 2005; Wohlwill, 1970). For convenience, however, we will use the phrase aging independently of the respective study’s sampling design in this article. The lack of longitudinal studies on sleep in adulthood does not allow for the identification of ‘real’ age-related changes of sleep physiology. Hence, we emphasize that longitudinal study designs are essential to study aging and to identify the lead-lag relations between age-related alterations in brain structure and function (Hofer and Sliwinski, 2001; Li and Schmiedek, 2002; Lindenberger, 2014; Raz and Lindenberger,2011).

#### Box 1. Sleep electrophysiology at a glance

In most animal species, three different and discrete states of vigilance can be differentiated: wakefulness, REM (rapid eye movement), and NREM (non-rapid eye movement) sleep (Vassalli and Dijk, 2009). REM and NREM sleep can be clearly distinguished based on electrophysiological characteristics (Aserinsky and Kleitman, 1953; Dement and Wolpert, 1958; Feinberg and Evarts, 1969; Loomis et al., 1935, 1962). REM sleep is marked by the occurrence of phasic irregular and rapid eye movements, muscle atony, and desynchronized wake-like electroencephalographic activity. NREM sleep, by contrast, is characterized by synchronous, low-frequency, high-amplitude EEG oscillations (Iber et al., 2007). NREM sleep can further be divided into the three sub-stages N1, N2 and N3 (also SWS; Iber et al., 2007; Rechtschaffen and Kales, 1968). These NREM sub-stages form a continuum of increasing arousal threshold and sleep depth (Carskadon and Dement, 2011). At the same time, they serve as a proxy for the dominance of different sleep-specific oscillatory brain signals: Lighter NREM sleep (stage 2 or N2 sleep) is defined by the occurrence of discrete sleep spindles (i.e., rhythmically waxing and waning oscillatory events with a typical frequency of 11–16 Hz), and K-complexes (i.e., phasic negative high-amplitude EEG deflections that are followed by a positive EEG component). With deeper sleep, the number and power of EEG slow waves increases. This predominance of slow EEG waves defines the presence of SWS (stage 3 or N3 sleep).

### Challenge 1: Ambiguous sleep stage definitions across age groups

*Standardized visual scoring of sleep stages is regarded as the ‘gold-standard’ of sleep research. Defined sleep stages are thereby considered sensitive indicators of specific physiological events. However, current sleep stage definitions may not capture the same physiological processes across age groups. We suggest that genuine sleep analysis – in line with similar views from animal research – should consider the continuous nature of NREM sleep. A focus on the presence of predefined electrophysiological events, like the occurrence or coupling of slow oscillations and sleep spindles, during **both** stage 2 sleep and SWS is required*.

#### Pitfalls of visually scoring SWS

In human research, the classification of sleep stages (cf. Box 1) is based on widely accepted standardized rules that are used to score sleep by means of EEG activity, muscle tone, and eye movements (Iber et al., 2007; Rechtschaffen and Kales, 1968). SWS, as defined by the *American Academy for Sleep Medicine* (AASM; Iber et al., 2007), requires the presence of 0.5–2 Hz waves with a minimum amplitude of 75 μV covering more than 20 % of a sleep time segment. Already in 1982, Webb noted that the frequently reported striking age-related reduction in SWS likely results from the use of a fixed amplitude threshold to define slow waves (Webb, 1982; Webb and Dreblow, 1982; cf. Figure 1A, Table 1). As also acknowledged by the AASM (Silber et al., 2007), reliably measuring SWS using an amplitude criterion below 75 μV is possible. Still, the current version of the manual for sleep scoring that defines the prevailing gold-standard, relies on a minimum amplitude criterion of 75 µV. Crucially, the age-related reduction in slow wave (< 4 Hz) and slow oscillation (< 1 Hz) amplitudes is one of the most consistent findings in age-comparative sleep studies (Carrier et al., 2011; Dubé et al., 2015; Mander et al., 2017; Muehlroth et al., 2019b; Ujma et al., 2019). Typically, SWS is easily identifiable in younger adults because slow waves exceed the required amplitude threshold. By contrast, the identification of true SWS in older adults is challenging: Due to the reduction of slow wave amplitudes below 75 μV (cf. Figure 1B and C, Table 2), time segments can often not be scored as SWS but are rather assigned to stage 2 sleep – although all other characteristics of the sleep epoch suggest the presence of SWS (i.e., the predominance of low-frequency EEG waves but absence of eye movements along with reduced muscle tone; cf. Figure 2).

**Figure 1.**
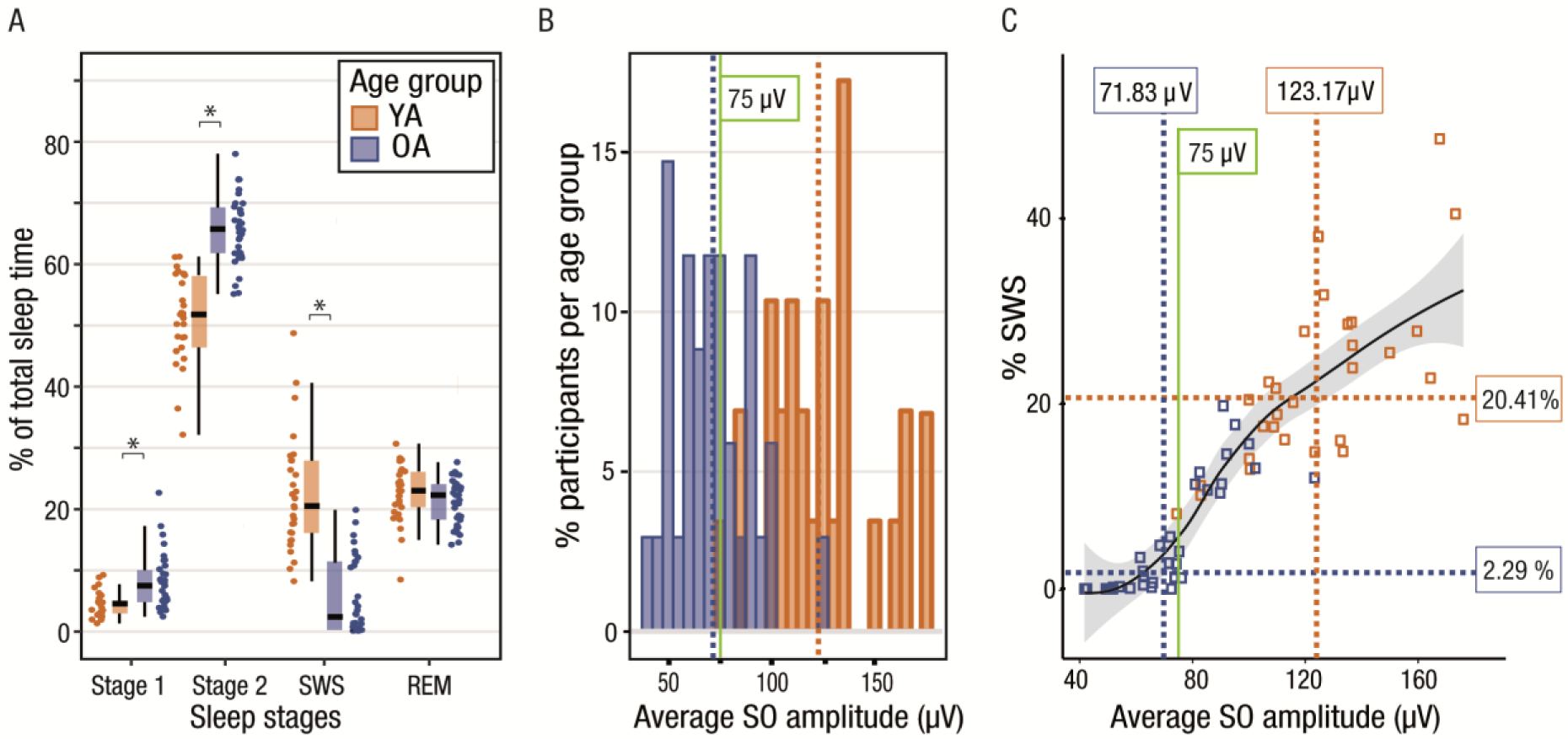
Visual sleep scoring in younger and older adults. (A) Distribution of visually scored sleep stages in younger (orange) and older (blue) adults. In older adults, the proportion of SWS is significantly reduced, whereas lighter stage 2 sleep is increased. Note the bimodal distribution of SWS in older adults with some participants displaying levels of SWS comparable to younger adults, and others clustering around 0 %. Asterisks mark *p*-values <.001 derived from non-parametric Mann-Whitney *U* tests comparing sleep parameters between younger and older adults (cf. Table 1). (B) Distribution of the average slow oscillation amplitude for all younger (orange) and older adults (blue). Age group medians are inserted as dashed lines. The median slow oscillation amplitude in older adults lies below the amplitude criterion of 75 μV (green line). (C) The proportion of visually scored SWS scales with the average slow oscillation amplitude (respective age group medians marked by dashed lines). Older adults falling below the 75 μV criterion (inserted in green) show little or no SWS. The smoothed local regression line (in black, fitted using LOESS) and the corresponding standarderror (grey shading) further illustrate the association. SO: slow oscillation.

**Figure 2.**
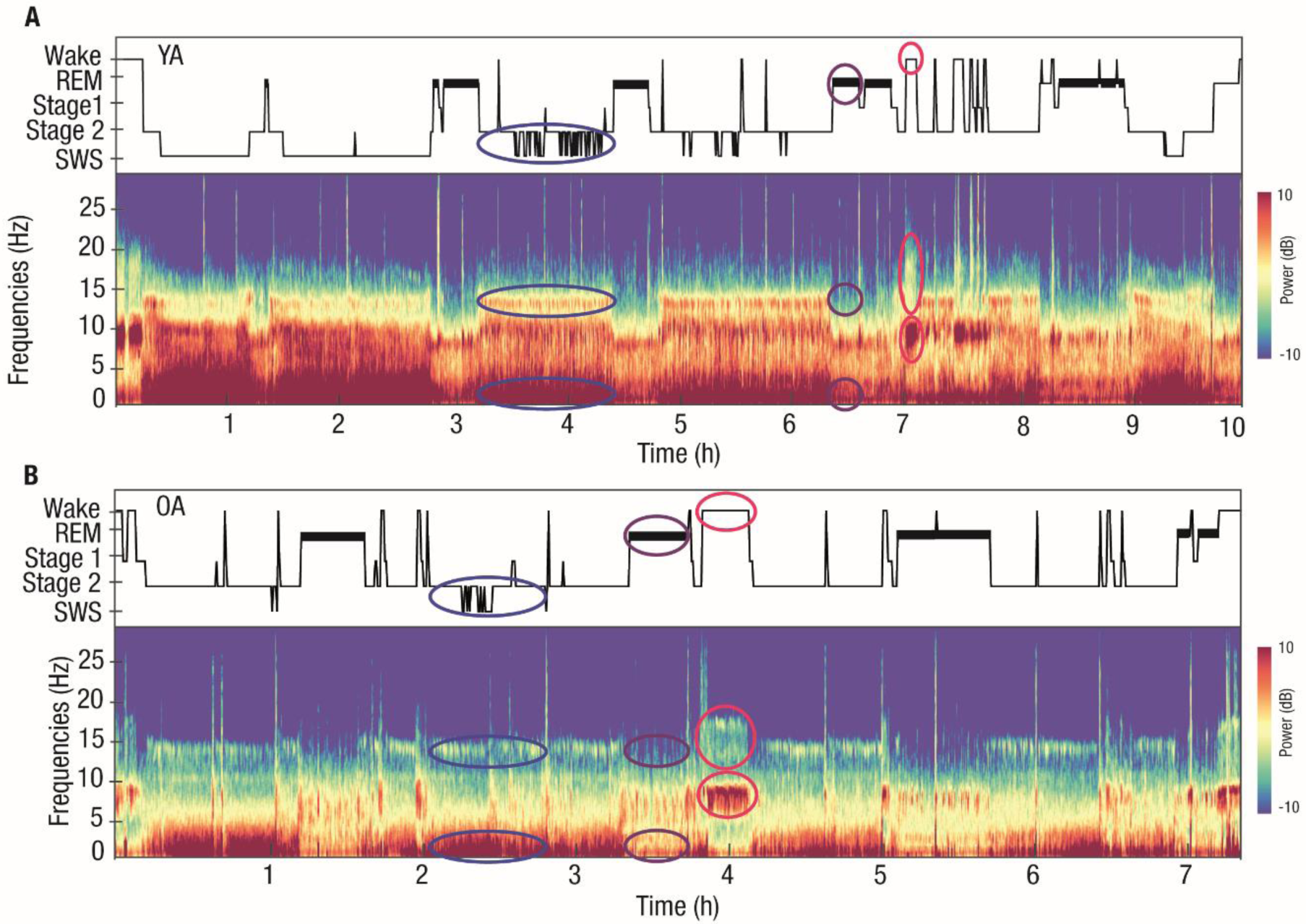
Whole-night spectral EEG characteristics. Hypnogram (upper panels) from one exemplary younger (A; female, 24.63 years) and older (B; female, 71.55 years) adult with the corresponding whole-night multitaper spectral EEG data at electrode Cz (lower panels). Both younger and older adults show a similarly increased low-frequency power during periods of NREM sleep (some examples highlighted by blue ellipses) along with a distinct increase in power in the fast spindle frequency range (approx. 12.5–16 Hz) that disappears during REM sleep (violet ellipses). Periods of wake are characterized by an additional strong enhancement of alpha (8–12 Hz) and high frequency power (red ellipses). Although the older adult only has a small amount of visually scored SWS, the spectrogram clearly shows periods of enhanced low frequency and spindle power – in line with the canonical SWS characteristics found in younger adults. No clear distinction between stage 2 and SWS is observed on the basis of spectral EEG characteristics. Note the differences in absolute sleep duration (x-axis). YA: younger adult; REM: rapid eye movement sleep; SWS: slow-wave sleep; OA: older adult.

**Table 1.** Age differences in sleep architecture

| | Younger Adults | Older adults | | | Cohen's $d$ |
| --- | --- | --- | --- | --- | --- |
| | $Md$ [1st qrt; 3rd qrt] | $Md$ [1st qrt; 3rd qrt] | $Z$ | $p$ | [95 % CI] |
| TST (min) | 456.50 [430.25; 496.75] | 425.50 [385.06; 459.88] | 2.48 | .013 | 0.62 [0.1; 1.15] |
| WASO (min) | 6.75 [3.00; 11.50] | 21.25 [12.06; 49.81] | -4.23 | <.001 | -1 [-1.54; -0.45] |
| Stage 1 (%) | 4.46 [2.84; 4.82] | 7.39 [4.71; 9.90] | -3.99 | <.001 | -1.02 [-1.57; -0.48] |
| Stage 2 (%) | 51.68 [46.29; 58.10] | 65.62 [61.71; 69.14] | -6.23 | <.001 | -2.95 [-2.96; -1.63] |
| SWS (%) | 20.41 [16.01; 27.83] | 2.30 [0.16; 11.15] | 6.03 | <.001 | 2.18 [1.53; 2.83] |
| REM (%) | 22.91 [20.22; 26.07] | 22.19 [18.22; 24.00] | 1.46 | .144 | 0.31 [-0.20; 0.83] |

*Note.* $Z$ and $p$ -values were derived from non-parametric Mann-Whitney $U$ tests comparing sleep parameters between younger and older adults. Sleep stage percentages are calculated as proportions of TST. CI: confidence interval; qrt: quartile; TST: total sleep time; WASO: wake after sleep onset (i.e., time awake between sleep onset and final morning awakening); SWS: slow-wave sleep; REM: rapid eye movement sleep.

**Table 2.**
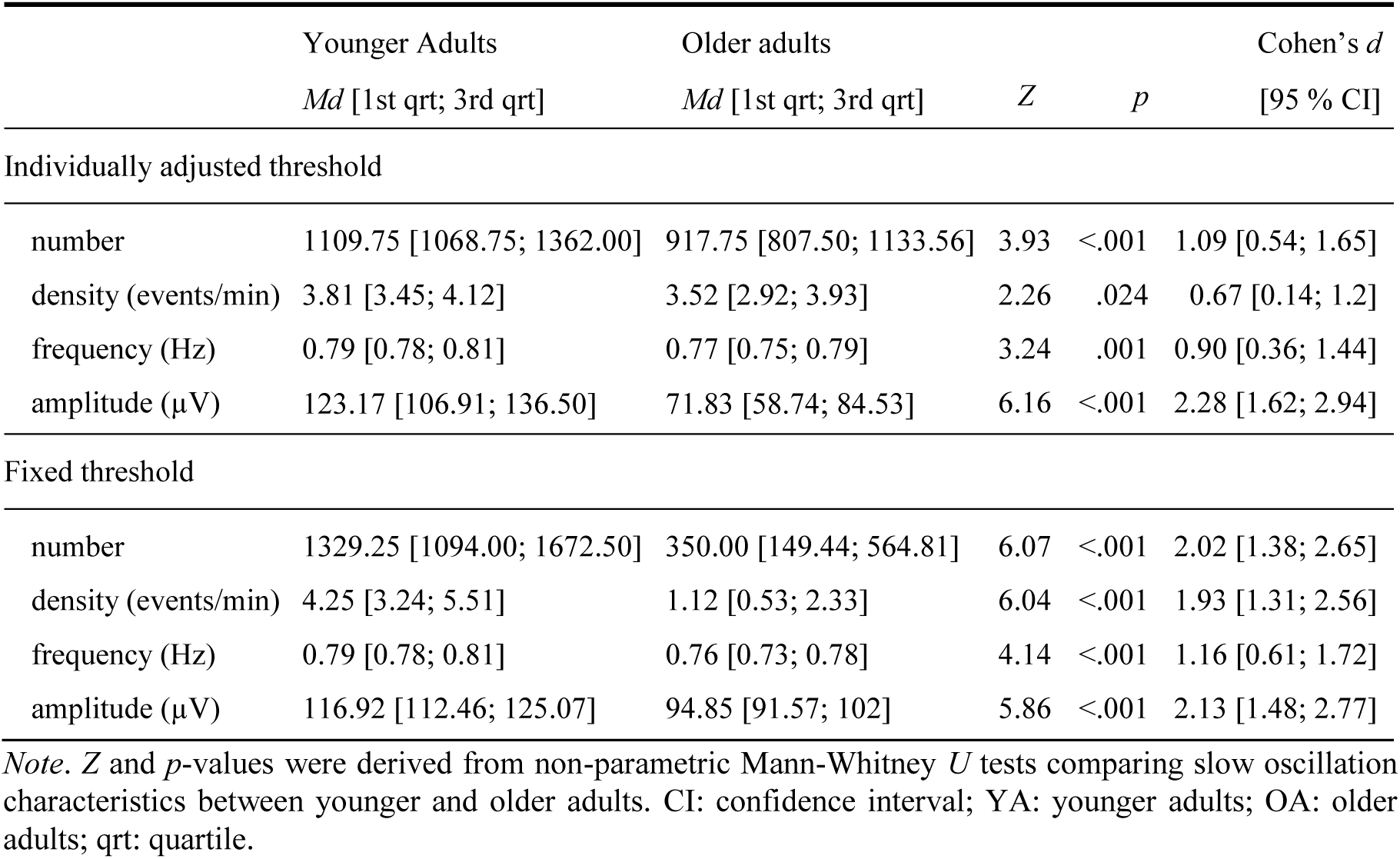
Individually adjusted vs. fixed threshold settings for slow oscillation detection

|  | Younger Adults<br>*Md* [1st qrt; 3rd qrt] | Older adults<br>*Md* [1st qrt; 3rd qrt] | *Z* | *p* | Cohen's *d*<br>[95 % CI] |
| --- | --- | --- | --- | --- | --- |
| Individually adjusted threshold |  |  |  |  |  |
| number | 1109.75 [1068.75; 1362.00] | 917.75 [807.50; 1133.56] | 3.93 | <.001 | 1.09 [0.54; 1.65] |
| density (events/min) | 3.81 [3.45; 4.12] | 3.52 [2.92; 3.93] | 2.26 | .024 | 0.67 [0.14; 1.2] |
| frequency (Hz) | 0.79 [0.78; 0.81] | 0.77 [0.75; 0.79] | 3.24 | .001 | 0.90 [0.36; 1.44] |
| amplitude (μV) | 123.17 [106.91; 136.50] | 71.83 [58.74; 84.53] | 6.16 | <.001 | 2.28 [1.62; 2.94] |
| Fixed threshold |  |  |  |  |  |
| number | 1329.25 [1094.00; 1672.50] | 350.00 [149.44; 564.81] | 6.07 | <.001 | 2.02 [1.38; 2.65] |
| density (events/min) | 4.25 [3.24; 5.51] | 1.12 [0.53; 2.33] | 6.04 | <.001 | 1.93 [1.31; 2.56] |
| frequency (Hz) | 0.79 [0.78; 0.81] | 0.76 [0.73; 0.78] | 4.14 | <.001 | 1.16 [0.61; 1.72] |
| amplitude (μV) | 116.92 [112.46; 125.07] | 94.85 [91.57; 102] | 5.86 | <.001 | 2.13 [1.48; 2.77] |

*Note.* *Z* and *p*-values were derived from non-parametric Mann-Whitney *U* tests comparing slow oscillation characteristics between younger and older adults. CI: confidence interval; YA: younger adults; OA: older adults; qrt: quartile.

Re-analyzing our own previously published data from 29 healthy younger (range: 19–28 years, *M*_age_ = 23.55 years, *SD*_age_ = 2.58; 16 females) and 34 healthy older adults (range: 63–74 years, *M*_age_ = 68.85 years, *SD*_age_ = 3.11, 15 females), we demonstrate the effects of fixed amplitude thresholds for visual sleep scoring in age-comparative studies. Participants’ nighttime sleep was monitored using ambulatory polysomnography (PSG) before and after completing an associative learning task (cf. Supplementary Methods; Muehlroth et al., 2019a,b). Subjective sleep quality was assessed using the Pittsburgh Sleep Quality Index (PSQI; Buysse et al., 1989) and comparable in both age groups (*Z* = 1.03, *p* = .305, *Md*_YA_= 4.5 [3.0; 6.25], *Md*_OA_= 4.0 [3.0; 5.0]). We show that, overall, in our dataset the extent of age-related reductions in visually scored SWS matched the increase in stage 2 sleep in older adults (Cohen’s *d*_stage 2_ = –2.3 [–2.96; –1.63]; Cohen’s *d* _SWS_ = 2.18 [1.53; 2.83]; Table 1). Visually scoring the data from older adults, a reduction – and in some participants even ‘absence’ – of SWS became evident in comparison to younger adults (Table 1, Figure 1A). Closely inspecting the sleep architecture of older adults in our sample, it became evident that the older age group was divided into subjects not exhibiting any or only few SWS, and subjects showing a more ‘youth-like’ proportion of SWS (Figure 1A). In part, this bisection of older subjects might be due to inter-individual differences in the amplitude of slow oscillations, with half of the older adults showing slow oscillation amplitudes below the proposed 75 μV threshold on average (*Md*_OA_ = 71.83 μV; Figure 1B). For older adults, falling above or below this threshold divided the sample into individuals with and without SWS according to the standard scoring system (Figure 1C). None of the younger adults undercut an average slow oscillation amplitude threshold of 75 μV (*Md*_YA_ = 123.17 μV) and, accordingly, SWS could be effectively scored in this age group (Figure 1B and C).

Taken together, it appears that due to age-related reductions in slow wave amplitudes, the fixed amplitude criterion of 75 μV for scoring SWS results in a biased estimate of SWS presence in older populations. Descriptively providing standardized estimates of sleep architecture is important to ensure the comparability and validity of studies. In studies with heterogeneous samples, though, researchers should acknowledge that such derived sleep stage estimates may be severely biased. As Rechtschaffen and Kales themselves put it in 1968 when establishing the standardized scheme for visual sleep scoring: *“Even among human subjects, however, there are some individuals or groups whose polygraph recordings may require further description or elaboration than that provided by the stages proposed here”* (Rechtschaffen and Kales, 1968).

#### The need to acknowledge the continuous nature of NREM sleep

The potential bias of current sleep stage definitions across age groups has direct theoretical and methodological implications. These affect the way we should conceive and treat NREM sleep and its sub-stages in our theories and analyses. The transition between stage 1, stage 2, and SWS happens continuously with slow waves increasing in amplitude and abundance as NREM sleep gets deeper. On the contrary, sleep spindles show a reversed pattern with predominance during lighter stage 2 NREM sleep (Prerau et al., 2017). Hence, stage 2 and SWS are often treated as proxies for the dominance of different sleep-specific oscillatory brain signals (Genzel et al., 2014; Hobson, 1968). Although generally accepted in human sleep research, the distinction between sleep stage 2 and SWS based on a certain amplitude and time criterion (Iber et al., 2007) is not without problems (Genzel et al., 2014; Lacroix et al., 2018; Webb, 1982; Silber et al., 2007). Challenges include the handling of potentially biased sleep stage estimates in different age groups and the comparability of estimates between human and animal research (e.g., Genzel et al., 2014; Lacroix et al., 2018).

As discussed above, the current criteria to define the presence of SWS along with great inter-individual variation in slow wave amplitudes may severely bias the assignment of a specific sleep segment to stage 2 sleep or SWS. Between the ages of 5 and 90, the proportion of NREM sleep decreases only little (Ohayon et al., 2004). The composition of visually scored NREM sleep itself, however, changes drastically across the lifespan. This might at least partly stem from the fact that time segments ‘equivalent’ to SWS may be scored as stage 2 sleep because the amplitude criterion of 75 μV is not met and result in two major problems: First, the assignment of SWS-like time segments to stage 2 sleep in older adults blurs the boundary between the two stages. As a consequence, sleep stages can no longer be regarded as a direct reflection or proxy of underlying neural and physiological processes (Hobson, 1968). Second, due to skewed distributions and reduced variance, any correlational analysis relying on measures of SWS is biased, or even prevented, in samples of older adults – especially in samples of older adults with a large percentage of ‘no-SWS’ participants. As a result, a clear-cut association between a given physiological state – identified via a corresponding sleep stage – and an outcome measure, like the ability to maintain memories across sleep (e.g., Diekelmann and Born, 2010; Rasch and Born, 2013), is hardly tangible in age-comparative studies. If the definitions used to identify a given sleep stage do not capture the same phenomenon (e.g., the presence of slow oscillations) to the same degree across age groups, age differences in associations with outcome measures can hardly capture the intended mechanism. For example, the observation that memory performance is more consistently associated with SWS in younger than in older adults, could be taken as evidence that slow oscillations forfeit their functionality for memory processing during aging (e.g., Baran et al., 2016; Scullin, 2013; Sonni and Spencer, 2015). However, if slow oscillations are present in older adults but just attributed to a different sleep stage, the statistics might be correct, but the functional interpretation would be different. In this case, slow oscillations found in a different sleep stage may continue to contribute to the consolidation of memories even in old age – a matter of fact only revealed when focusing on the oscillatory phenomena themselves (e.g., Helfrich et al., 2018; Mander et al., 2013; Muehlroth et al., 2019b; Varga et al., 2016; Westerberg et al., 2012).

Given the described challenges a NREM sleep subdivision bears, it may be hardly surprising that research in other species refrains from applying such a sub-classification (but see Lacroix et al., 2018, for a recent attempt for ‘human-like’ sub-classification of NREM sleep in rodents). The term ‘SWS’ is typically used to describe the whole continuum of NREM sleep, which has evoked pivotal misunderstandings when comparing human and animal research (Genzel et al., 2014). This conception of NREM sleep may arise from the prevailing sleep scoring procedure in rodents that can easily distinguish between REM and NREM sleep on the basis of movements and spectral parameters of electrical neuronal activity during sleep, but less so between NREM sub-stages (Bagur et al., 2018; Lacroix et al., 2018). In humans, contemplating whole-night multitaper spectral EEG data irrespective of participants’ age, the distinct natures of REM and NREM sleep becomes evident, whereas differences between SWS and stage 2 sleep are more subtle as they transition gradually (Figure 2; see Prerau et al., 2017 for a detailed description of the spectral dynamics of the whole-night multitaper spectogram). Crucially, this does not signify that NREM sleep should be regarded as a uniform state, rather that the continuous nature of NREM sub-stages must be acknowledged, which makes a clear-cut distinction and definition of stage 2 and SWS problematic, if not impossible.

To conclude, an explicit distinction between stage 2 and SWS bears not only methodological but also theoretical difficulties. A coherent methodology to study the complex and dynamic nature of sleep not only within, but also between species is required to enable informative study comparisons (cf. Prerau et al., 2007). Research on the causes and consequences of inter-individual and age-related differences in sleep physiology should thus transcend stage-based analyses and rather consider the actual *exploranda*, ie., the underlying neurophysiological processes (for instance, the occurrence of rhythmic thalamocortical sleep spindles) present during **both** stage 2 sleep and SWS.

##### Box 2. Guidance for the interpretation of frequently reported oscillatory characteristics

*Spectral power*. The spectral power in predefined frequency bands is believed to mirror the presence of rhythmic neural activity, i.e., periodic alternations in the excitability of neuronal populations (Buzsáki, 2006; Buzsáki et al., 2016; Cohen, 2014). Yet, conventional estimates of spectral power reflect a compound of rhythmic and arrhythmic activity and, within their rhythmic component, index a combination of the duration and the amplitude of oscillations within a specific frequency range (Cohen, 2014; Kosciessa et al., in press).

*Amplitude*. The amplitude of an oscillation reflects the degree of synchronized neuronal firing: Small amplitudes relate to more local synchronous firing, whereas large amplitudes reflect more global synchronous changes in membrane potentials (Buzsáki, 2006; Nir et al., 2011). The basis for this cortical synchronization is the integrity of cortical areas involved in the generation and propagation of a certain oscillation (Dubé et al., 2015; Saletin et al., 2013). In aging, cortical thinning and atrophy may diminish the scalp-detectable range of amplitudes. Age differences in oscillatory amplitudes may, on the one hand, reflect a minimized extent of synchronous neuronal firing as direct consequence of structural brain changes in the course of aging (Dubé et al., 2015). On the other hand, anatomical alterations of the brain and skull may influence the conduction of the EEG signal and thus attenuate the amplitude of an oscillation measured on the scalp (Leissner et al., 1970; Segalowitz and Davies, 2004). Besides, reduced slow wave amplitudes in old age may arise from alterations in homeostatic sleep pressure, which is typically reduced as humans age (Dijk et al., 2000; Esser et al., 2007; Vyazovskiy et al., 2009).

*Frequency*. In general, the inherent frequency of an oscillation may index the efficiency and speed of neural information transfer (Dubois et al., 2008; Grandy et al., 2013; Klimesch, 1999; Posthuma et al., 2001). The typical developmental acceleration of various oscillatory components including sleep spindles (Campbell and Feinberg, 2016; Purcell et al., 2017) is believed to depend on the differentiation of thalamocortical feedback loops due to maturation, optimization of synaptic connections, and neuronal myelination (Campbell and Feinberg, 2016; Clawson et al., 2016; Dubois et al., 2008; Klimesch, 1999; Piantoni et al., 2013). Maintained or even increased frequency of fast spindles in old age (Crowley et al., 2002; Fogel et al., 2017; Nicolas et al., 2001; but see Martin et al., 2013) may indicate a largely preserved functionality of thalamocortical cells during aging. In contrast, the consistently reported slowing and flattening of slow oscillations in old age (Carrier et al., 2011; Dubé et al., 2015; Mander et al., 2017) may be the consequence of reduced synaptic load and homeostatic sleep pressure in old age and indicative of decelerated and less synchronous switches between phases of neuronal excitability and inhibition (Bersagliere and Achermann, 2010; Carrier et al., 2011; Dijk et al., 2000; Dubé et al., 2015; Fogel et al., 2012; Vyazovskiy et al., 2009

### Challenge 2: Multiple ways to describe sleep physiology

*The multitude of indicators and available algorithms used to describe sleep-specific neural activity hampers the comparability of results between studies. We argue that the validity and value of age-comparative sleep research depends greatly on the precise definition of indicators that reflect a neural process of interest as directly as possible. In our view, indicators of sleep micro-structure should always be preferred as they more closely mirror the dynamics and characteristics of sleep-specific neural activity*.

Everyone engaging in sleep research will soon realize the multitude and inconsistency of indicators available to describe neurophysiological activity during sleep (Supplementary Figure 1). Comparing different measurement occasions (Brandmaier et al., 2018) and diverse indicators of slow oscillatory activity during NREM sleep in our own data set (namely % SWS, relative SWA, and the number, density, frequency, and amplitude of detected slow oscillations), we noticed a generally poor agreement between the different variables both across and within age groups (*ICC* = 0.201 [–.15; .47], *F*(62,310) = 1.25, *p* = .113; *ICC*_YA_ = 0.13 [–.48; .54], *F*_YA_(28,140) = 1.15, *p*_YA_ = .295; *ICC*_OA_ = 0.09 [–.49; .49], *F*_OA_(33,165) = 1.1, *p*_OA_ = .342; cf. Supplementary Table 1). In contrast, the test-retest reliability of the same variable between two consecutively recorded nights was good to excellent (all *ICC ≥* .78, all *F ≥* 4.61, all *p* < .001; cf. Supplementary Table 1). A moderate test-retest reliability of relative SWA in the group of older adults was the only exception (*ICC* = .68 [.33; .85], *F*(33,33) = 3.69, *p* < .001). Critically, all participants engaged in an extensive associative learning task between the two recorded nights (cf. Muehlroth et al., 2019a,b). As demanding learning is known to influence indicators of spindle and slow oscillatory activity (e.g., Gais et al., 2002; Mölle et al., 2011, 2009), the reported estimates should even be considered as the lower bounder of reliability. Taken together, even though sleep physiology can be very reliably measured by applying common analytic approaches, the great disparity of available indicators calls for a thorough consideration of the way we describe sleep physiology in our analyses.

In general, the validity of indicators to quantify neural activity during sleep scales with their resolution and precision: Age differences in sleep are best captured by describing the dynamics and features of neural activity in as much detail and as directly as possible. Using machine learning on 3500 sleep recordings covering the ages 18 to 80, Sun and colleagues (2019) recently demonstrated that oscillatory dynamics during sleep – like fluctuations in spectral power – (i.e., the *micro-structure* of sleep) can reliably predict participants’ age (mean absolute deviation 7.8 years; see Sun et al., 2019). Global sleep stage composition (i.e., the *macro-structure* of sleep), in contrast, turned out to be a less appropriate predictor of chronological age (mean absolute deviation 23.3 years; Sun et al., 2019). Sleep is known to be a dynamic process (Prerau et al., 2017; Spiess et al., 2018; Yetton et al., 2018). The use of sleep stage estimates mismatches our current theoretical understanding of sleep and strongly reduces resolution of the data (Conte and Ficca 2013; Prerau et al., 2017). Altogether, we believe that the analysis of sleep EEG data should transcend beyond the report of sleep stages and global spectral features and focus on the micro-structure of sleep.

To avoid using the proportion of SWS as a proxy for the presence of slow waves, analyses increasingly report on *slow-wave activity* (SWA). SWA measures the spectral delta power (i.e., 0.5–4.5 Hz) via a transformation of the data in the frequency domain (e.g., a Fast Fourier Transformation, FFT). In conjunction with a normalization procedure – e.g., dividing the respective frequency band by the total power of the whole spectrum – the influence of possible confounds such as frequency-unspecific inter-individual differences in EEG amplitudes can be minimized (see, e.g., Dannhauer et al., 2011; Frodl et al., 2001; Leissner et al., 1970; Segalowitz and Davies, 2004; Werkle-Bergner et al., 2009, for arguments on how inter-individual differences in anatomical properties of the brain and skull influence the EEG signal). A normalized measure of SWA increases the comparability across diverse study populations. However, spectral power, as conventionally measured with FFT, is a compound of rhythmic processes and the arrhythmic 1/f background spectrum that, crucially, is flattened in older compared to younger adults (Voytek et al., 2015; Supplementary Figure 2; see Haller et al., 2018; Kosciessa et al., in press; Wen and Liu, 2016; Whitten et al., 2011 for approaches to separate rhythmic activity from the arrhythmic background spectrum). Moreover, as spectral power conflates both duration and amplitude of rhythmic neural activity (e.g., Kosciessa et al., in press), an interpretation of SWA in terms of abundance or strength of slow waves remains difficult (Amzica and Steriade, 1998; Mensen et al., 2016). *“Thus it is essential to strive toward a characterization of sleep EEG oscillations that faithfully represent the underlying data, allowing us to apply the wealth of knowledge we have gained about the continuum of the underlying neurophysiological mechanisms to the interpretation of the EEG”* (Prerau et al., 2017, p. 63).

Technological developments over the past decades are increasingly facilitating analysis of specific features and characteristics of neural events during sleep (Mensen et al., 2016; Mölle et al., 2002; Prerau et al., 2017). Such analyses can incorporate the amplitude, oscillatory frequency, the density of occurrence (i.e., number of events per time interval), and the slope of specific oscillatory events. These different indicators are often used interchangeably although they signify differential physiological properties and, accordingly, show differential age dependency (cf. Box 2, Supplementary Figure 1). If applied correctly, though, they hold great explanatory power and can provide more informed insights into the aging brain (Fogel et al., 2017; Sun et al., 2019; Ujma et al., 2019). Moreover, the possibility to identify individual oscillatory events also entails the opportunity to investigate how different oscillations and neural processes interact (Helfrich et al., 2018; Klinzing et al., 2016; Muehlroth et al., 2019b; Staresina et al., 2015). At the moment, due to the variety of algorithms available to study oscillatory brain signals and their interaction during sleep, these analyses are still time-consuming and between-study comparability of indicators can be challenging (Figure 3; cf. Warby et al., 2014). To tackle these problems, currently, several open source projects are under development that aim to harmonize and standardize analysis pipelines and parameters and make them publically available (e.g., Luna [http://zzz.bwh.harvard.edu/luna/], Sleep [Combrisson et al., 2017; http://visbrain.org/sleep.html], SPISOP [www.spisop.org], YASA [http://doi.org/10.5281/zenodo.3382284]; all code and sample data to reproduce this manuscript’s EEG analysis pipeline will be available on the Open Science Framework [https://osf.io/zur2b/]). Identifying and resolving these discrepancies – ideally by the use of common algorithms – will lay the grounds for a mechanistic understanding of the causes and consequences of age-related changes in sleep (Helfrich et al., 2018; Muehlroth et al.,2019b).

**Figure 3.**
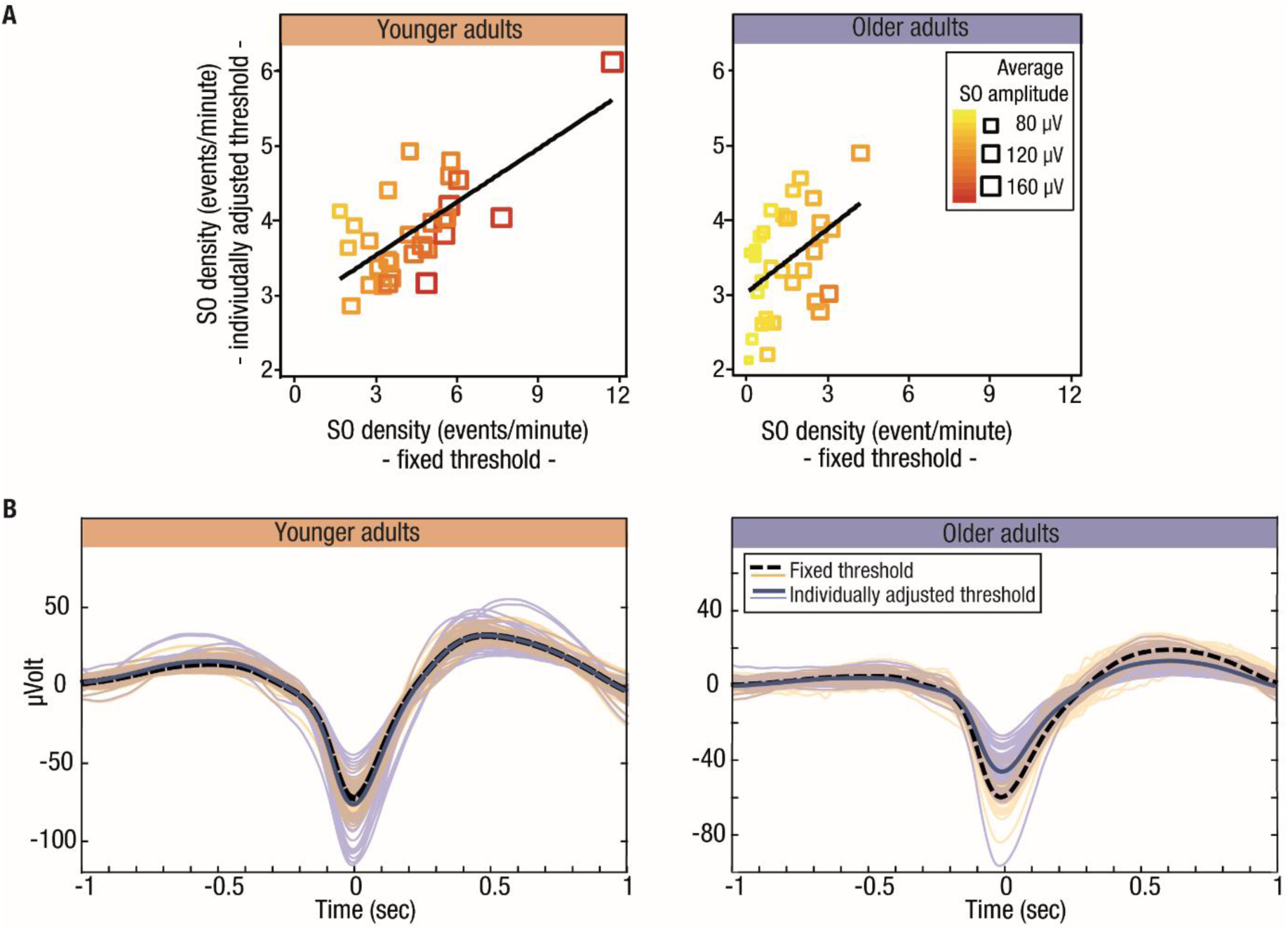
Comparison of fixed (75 μV) and individually adjusted slow oscillation detection thresholds. (A) Relation between slow oscillation density estimates derived by the use of individually adjusted vs. fixed detection thresholds. In younger adults (orange), the two detection thresholds produce comparable results. In older adults (blue), especially for individuals with small slow oscillation amplitudes (smaller square size), a fixed detection threshold results in an underestimation of slow oscillation density. (B) Individual average slow oscillations and age group averages of slow oscillations detected with individually adjusted and fixed amplitude thresholds. Left panel: In younger adults, both algorithms result in comparable slow oscillation detection. Right panel: In older adults, individually adjusted thresholds (blue) result in the detection of slow oscillations with smaller amplitudes compared to higher-amplitude fixed threshold detection (green). OA: older adults; SO: slow oscillation; YA: younger adults.

### Challenge 3: Amplitude reductions – towards algorithms with adjusted thresholds

*The benefits of a direct investigation of slow oscillatory events are constrained by a lacking consensus on the detection thresholds to be applied. We suggest that a fair assessment of true slow oscillatory activity requires the use of amplitude thresholds that are adjusted individually within participants*.

Studying rhythmic neural activity during sleep requires a reliable detection of oscillatory events themselves. Here, automated slow oscillation detection methods offer an efficient way to identify the presence and characteristics of individual slow oscillations (eg., Massimini et al., 2004; Mensen et al., 2016; Mölle et al., 2002; Nir et al., 2011). Typically, oscillation detection algorithms rely on a bandpass-filtered EEG signal within a theoretically implied frequency range (eg., 0.5–1 Hz for slow oscillations, see Challenge 4 for details on the definition of frequency ranges). The presence of oscillatory components is then inferred by applying certain amplitude and peak detection criteria (cf. Supplementary Methods). Amplitude criteria can either be based on fixed thresholds or on variable thresholds adjusted within individuals (e.g., Massimini et al., 2004; Mölle et al., 2009, 2002; Staresina et al., 2015). In analogy to the visual scoring of SWS, many algorithms rely on a criterion of 75 μV (Carrier et al., 2011; Dubé et al., 2015; Piantoni et al., 2013) or even higher amplitudes (Heib et al., 2013; Massimini et al., 2004) for the detection of slow oscillations during NREM sleep. But an increasing number of studies opt for thresholds adjusted individually for each participant (e.g., Klinzing et al., 2016; Mölle et al., 2011; Ngo et al., 2015). As yet, there is neither a gold-standard nor consensus on the most suitable method nor on the threshold settings to be applied when identifying the presence of oscillatory events (Mensen et al., 2016; Ujma et al., 2019). However, the choice of detection thresholds has profound effects on the analysis of age differences.

Irrespective of the applied detection threshold (i.e., a fixed threshold of 75 μV vs. an individually adjusted threshold of 1.25 times the mean amplitude of all putative slow oscillations; cf. Supplementary Methods), in the older adults in our data, the number, density (i.e., the number of slow oscillations per minute of NREM sleep), amplitude, and frequency of slow oscillations were all reduced (Table 2). Nevertheless, compared to an individually adjusted amplitude criterion, the extent of age effects was augmented when using a fixed detection threshold of 75 μV (non-overlapping confidence intervals for Cohen’s *d* of slow oscillation density, cf. Table 2). Using individually adjusted thresholds, as displayed in Figure 1B, in older adults, the average slow oscillation amplitude fell below the 75 μV criterion in 61.77 % of the subjects (21 out of 34 compared to only one younger subject [3.45 %]). In other words, in older adults, a large number of potential slow oscillations will not be detected in case of a fixed amplitude criterion of 75 μV as the detection is biased towards higher-amplitude oscillations (Figure 3B). As a consequence, slow oscillation density estimates that are derived using individually adjusted vs. fixed detection thresholds are comparable in younger but not in older adults (*Z_YA_*= –.88, *p_YA_* = .38; *Z_OA_* = 6.08, *p_OA_* < .001; see Table 2; Figure 3A).

Critically, inter-individual differences in slow oscillation amplitudes might not only carry functional information but may be confounded by differences in homeostatic sleep regulation (Dijk et al., 2000), brain integrity (Dubé et al., 2015; Saletin et al., 2013), and skull thickness (Leissner et al., 1970; Frodl et al., 2001). Hence, an individually adjusted amplitude threshold that tags slow oscillations with the largest amplitudes within individuals may have advantages. Only when slow oscillations are selected independently of inter-individual differences in EEG amplitudes, do unbiased comparisons become possible. Persisting age differences in the amplitude, density, and morphology of slow oscillations can then be interpreted as valid age group differences. Compared to fixed amplitude detection criteria, especially in age-comparative settings, the use of individually adjusted amplitude criteria for detecting slow oscillations during NREM sleep provides a more realistic and at the same time less biased measure of true slow oscillatory activity.

### Challenge 4: Differential frequency shifts in sleep oscillations – towards individualized frequency ranges

*The exact frequency band in which an oscillatory event can be observed varies between individuals and is influenced by age. To prevent missing true oscillatory events during detection and mixing up functionally distinct oscillatory components, cut-off frequencies have to be determined carefully. We argue that individually determined frequency ranges are the prerequisite to derive meaningful age- and person-specific characteristics of oscillatory components*.

#### Contemplating frequency cut-offs for low-frequency oscillatory components

Age-related changes in the appearance of sleep oscillations do not only include alterations in amplitudes, but also entail shifts in the intrinsically expressed frequencies of the respective neural events. With respect to low-frequency components, a global slowing has consistently been reported (Carrier et al., 2011; Dubé et al., 2015; Mander et al., 2017) (Figure 4I, Box 2). These effects comprise both slow oscillations (< 1 Hz) as well as slow waves with higher frequencies (1–4 Hz). Although slow oscillations and slow waves constitute different entities on a theoretical level (Achermann and Borbély, 1997; Amzica and Steriade, 1998; Steriade et al., 1993), the applied cut-off frequencies defining low-frequency EEG components are often not much attended to.

**Figure 4.**
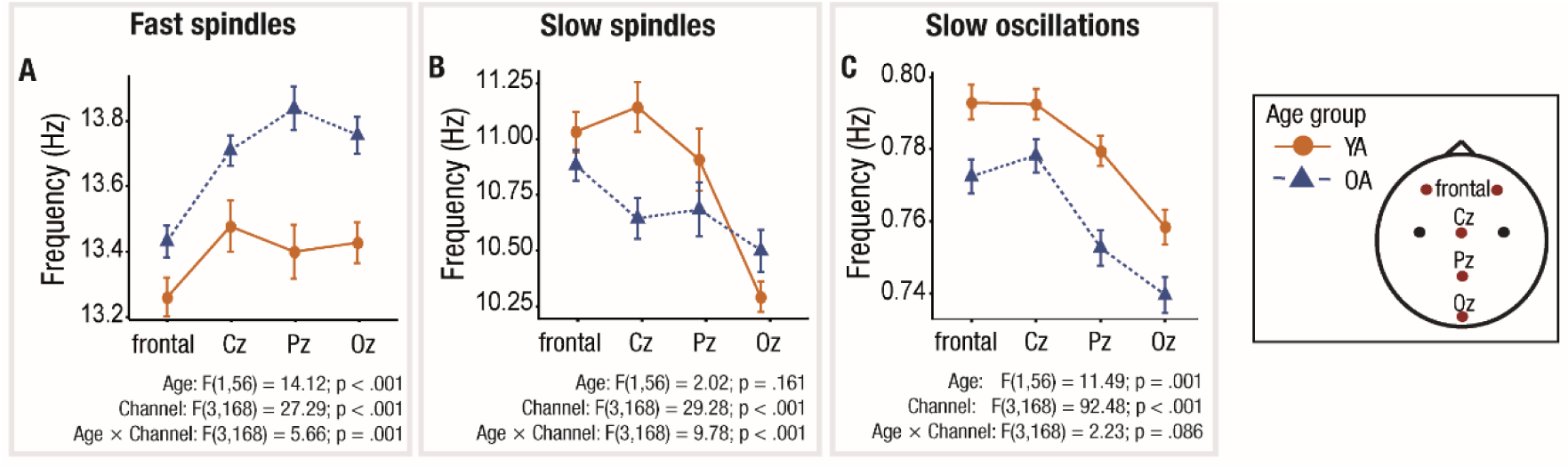
Topographical distribution of the average slow oscillation and spindle frequencies. (A) Overall increased fast spindle frequency in older (blue) compared to younger (orange) adults, with reduced effects at frontal channels. (B) Decreases in slow spindle frequency in older adults are only pronounced at Cz. (C) Uniform decrease in slow oscillation frequency in old age at all channels. Note. All slow oscillation and spindle characteristics were derived using individually adjusted amplitude thresholds. OA: older adults; SO: slow oscillation; YA: younger adults.

Empirically, the problem of identifying the exact frequency ranges for specific oscillatory components in individual participants can be solved by considering the NREM sleep power spectrum. Peaks in the power spectrum signify the presence of rhythmic neural activity at a given frequency (Aru et al., 2015; Kosciessa et al., in press; Whitten et al., 2011). When visually inspecting the age-specific power spectra during NREM sleep in our data, it became evident that the peak in low-frequency power was shifted to even lower frequencies in older adults (Supplementary Figure 2). This slowing in low frequencies was statistically reliable (average frequency peak: *t*(115) = 6.18, *p* < .001, *M*_YA_ = 0.73 Hz, *M*_OA_ = 0.53 Hz) and was also evident for discrete slow oscillation events (*Z* = 3.24, *p* = .001, *Md*_YA_= 0.79 Hz, *Md*_OA_ = 0.77 Hz; see Table 2, Figure 3B, Figure 4C). Although statistically detectable, in contrast to the dramatic reduction in slow oscillation amplitudes, inter-individual and age differences in slow oscillation frequencies were relatively small (*d*_frequency_ = 0.90 [0.36; 1.44]; *d*_amplitude_ = 2.28 [1.62; 2.94], non-overlapping confidence interval; cf. Table 2).

Taken together, it appears that the magnitude of the slowing of slow oscillations in old age does not reach an extent that would bias detection results when established frequency ranges (e.g., 0.5–1 Hz for slow oscillations) are utilized. Nonetheless, we emphasize the need to carefully consider, justify, and report the selected frequency range when publishing sleep research findings on older adults. Slow oscillations and ‘faster’ slow waves have been associated with differential homeostatic regulation (Bersagliere and Achermann, 2010; Werth et al., 1997) as well as with divergent roles in the context of cognitive pathology in aging (Lucey et al., 2019; Mander et al., 2015, 2016). Yet, detection algorithms often deploy frequency bands that comprise both oscillatory components (e.g., Carrier et al., 2011; Kurth et al., 2010). Finally, one should bear in mind that even when using adequate frequency cut-offs, filtering can induce signal distortions. The choice of filter type and filter parameters will differentially influence the signal and change detection results (Widmann et al., 2015). Together, we thus underscore the need for clear and coherent terminology along with appropriate parameter settings and detection criteria when reporting on age differences in low-frequency oscillatory activity.

#### Inter-individual (age) variation in fast and slow spindle frequency bands

In contrast to slow oscillations whose detection seems mostly dependent on appropriate amplitude settings, the detection of sleep spindles is heavily influenced by the selection of frequency ranges. In particular, the detection of spindle events is challenged by the presence of two spindle types. Slow (ca. 9–12.5 Hz) and fast spindles (ca. 12.5–16 Hz) differ in frequency, topography (Figure 4) (De Gennaro and Ferrara, 2003; Doran, 2003; Klinzing et al., 2016; Schabus et al., 2007), generating mechanisms (Ayoub et al., 2013; Schabus et al., 2007; Timofeev and Chauvette, 2013), and not least in their functionality (Barakat et al., 2011; Fogel and Smith, 2011; Mölle et al., 2011). Although the existence of two spindle types is now widely acknowledged, there is no general consensus on the exact definition of the frequency range in which to expect slow and fast spindles should be expected (Cox et al., 2017). Critically, spindles undergo a complex developmental shift and differentiation in their inherent frequency, power and topography (Campbell and Feinberg, 2016; Crowley et al., 2002; Nicolas et al., 2001; Purcell et al., 2017). This not only challenges research on younger adults (Cox et al., 2017), but also research that focuses on the entire lifespan.

In line with these observations, slow and fast spindle frequencies were differentially affected by age in our data. Average fast spindle frequency was increased in older adults (*F*(1,56) = 14.12, *p* < .001, *M*_YA_= 13.39 Hz, *M*_OA_ = 13.68 Hz) (Figure 4A, Box 2). In contrast, an age-related decrease in the average frequency of slow spindles was only pronounced at electrode Cz (age by channel interaction: *F*(3,168) = 9.78, *p* < .001, *Z*_Cz_ = 3.31, *p* < .001, *Md*_YA_ = 11.28 Hz, *Md*_OA_ = 10.61 Hz; all other |*Z| ≤* 1.41, all other *p ≥* .158, α-level adjusted to .0125; Figure 4B). The complexity of frequency shifts during aging is thus even enhanced by the topographical intricacy of the observed effects.

The multiple facets of age differences in sleep spindle appearance require the careful consideration of an age-adjusted definition of spindle frequency ranges. Based on individually identified peaks in the NREM sleep power spectrum at topographical locations where slow and fast spindles are expected (cf. Supplementary Figure 2), individually adjusted spindle frequency ranges can be defined – even if they might not match conventionally prescribed bands exactly (Adamczyk et al., 2015; Bódizs et al., 2009; Cox et al., 2017; Doppelmayr et al., 1998; Grandy et al., 2013; Klimesch, 1999; Ujma et al., 2015). Comparisons between spindle detection algorithms that use commonly predefined frequency ranges and those that adjust their frequency range definitions within individuals, provide a strikingly poor agreement with conventional (fixed frequency) algorithms. The latter have been shown to miss up to 25 % of fast and slow spindles (Adamczyk et al., 2015; Cox et al., 2017; Ujma et al., 2015). To allow for the investigation of differences in spindle characteristics, age-comparative research should ensure that the frequency bands applied to define slow and fast spindles are congruent with age- and person-specific spindle properties. Moreover, frequency definitions need to be broad enough to capture sleep spindles in their complexity and entirety. Independently of an individual’s age, person-specific definitions of spindle frequencies form the basis for these improvements (Adamczyk et al., 2015; Cox et al., 2017; Ujma et al., 2015).

### Challenge 5: Topographical disparities in age-related sleep changes

*Age-related alterations in sleep physiology do not show a spatially coherent pattern. We highlight that alterations in sleep oscillations that are measurable on the scalp do not necessarily reflect an altered initiation of underlying physiological sleep processes per se, but could rather represent age-related alterations in structural properties of the brain. Taking into account that intrinsic characteristics of sleep-specific oscillatory components may be well preserved during aging can help our understanding of the functionality of sleep in old age*.

Across the lifespan, the human brain undergoes complex structural alterations (Buckner, 2004; Cabeza et al., 2005; Fjell et al., 2013; Fjell and Walhovd, 2010; Raz et al., 2005), which are also reflected in the topography of the sleep EEG (Buchmann et al., 2011; Carrier et al., 2011; Kurth et al., 2010; Landolt and Borbély, 2001; Martin et al., 2013; Sprecher et al., 2016). From childhood to adolescence, along with cortical maturation in frontal brain regions, maximal SWA shifts from posterior to frontal brain regions (Kurth et al., 2010; Whitford et al., 2007). In old age, frontal brain regions are particularly prone to age-related structural white and gray matter loss (Giorgio et al., 2010; Raz and Rodrigue, 2006). As a consequence, age-related decline in spectral EEG power and amplitude is particularly pronounced over frontal areas (Carrier et al., 2011; Landolt and Borbély, 2001; Mander et al., 2017; Martin et al., 2013; Sprecher et al., 2016).

Topographical shifts in sleep parameters are diverse. For example, in our own data, the centroparietal prevalence of fast spindle density and amplitude in younger adults was shifted posterior in older adults, resulting in reduced or even absent age differences over parietal and occipital regions (Figure 5A and D; Supplementary Figure 2). In contrast, slow spindles and slow oscillations maintained their frontal dominance in old age. Whereas the amplitude and density of slow oscillations were globally and uniformly reduced in old age (Figure 5C and F), the density and amplitude of slow spindles showed most pronounced age-related reductions over frontal areas (Figure 5B and E). Thus, the extracted oscillatory event or indicator (i.e., slow spindle, fast spindle, slow oscillation), their derived characteristics (i.e., density, amplitude, and frequency), and not least an individual’s age interact and influence where, and to what extent, age differences in sleep can be observed.

**Figure 5.**
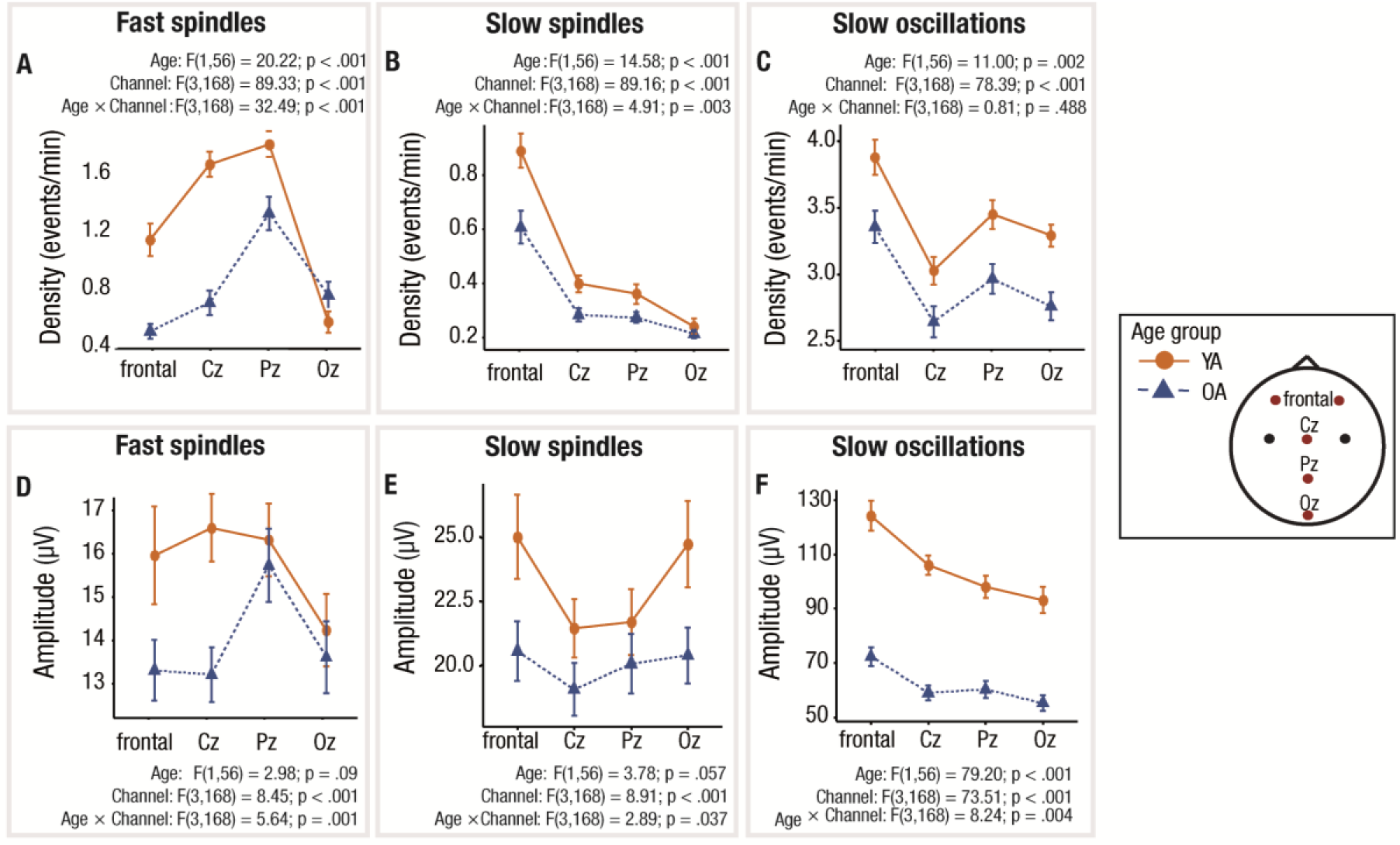
Topographical distribution of the average slow oscillation and spindle density and amplitude. (A) Centroparietal prevalence of fast spindles in younger adults (orange), compared to a parietal-occipital prevalence in older adults (blue). (B) Frontal prevalence in slow spindle density in both younger and older adults, as well as anterior to posterior attenuation of age differences. (C) Overall reduced slow oscillation density in older adults but preserved frontal dominance. (D) Preserved amplitude of parietal-occipital fast spindles in older adults. (E) Reduced slow spindle amplitude over centroparietal regions in younger adults, but more uniform amplitude modulation in older adults. (F) Overall reduced slow oscillation amplitude in older adults but anterior to posterior attenuation of age differences. Note. All slow oscillation and spindle characteristics were derived using individually adjusted amplitude thresholds. OA: older adults; SO: slow oscillation; YA: younger adults.

Crucially, the topographical heterogeneity of age effects detectable on the scalp level may be meaningful and indicative of alterations in underlying brain structure and cellular properties. To illustrate this effect, we tested for the association between gray matter volume and fast spindle density measured at different electrode sites (separate voxel-vise multiple regression models for each predictor; see Figure 6; cf. Supplementary Methods; Supplementary Figure 4, Supplementary Table 3). In line with the observed posterior preservation of fast spindles in old age (Figure 4A; Supplementary Figure 2), fast spindle density at Pz and Oz did not relate to gray matter volume. In contrast, fast spindle density measured frontally and at Cz, that was most reduced on older adults, was associated with structural integrity in frontal and temporal brain regions (Figure 6; Supplementary Figure 4, Supplementary Table 3). In comparison, gray matter clusters for the association with slow spindle and slow oscillation density were small and less consistent (Supplementary Figures 5 and 6, Supplementary Tables 4 and 5). Based on these analyses examples, it becomes clear that structural brain integrity does not globally shape the generation and propagation of all oscillatory components. However, one should note that chronological age, in general, was associated with global reductions in gray matter volume that were strongest in frontal brain regions (Figure 6; Supplementary Figure 3, Supplementary Table 2). The extent of structural brain alterations in aging can thus overshadow more focalized and potentially meaningful structure– function associations.

**Figure 6.**
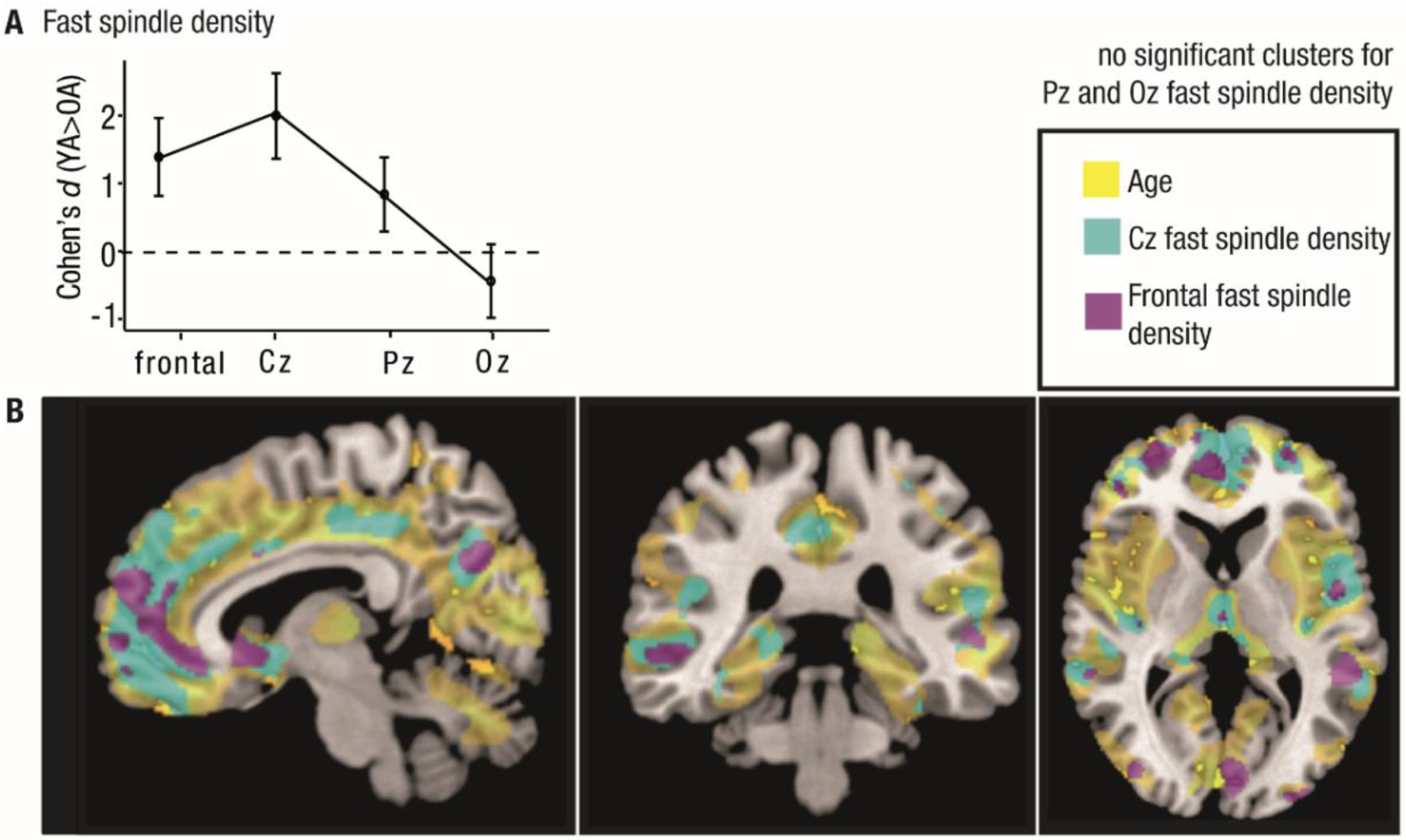
Age differences in fast spindle density mirror cortical integrity. (A) Cohen’s *d* of age differences in fast spindle density at different electrode sites. Confidence intervals are displayed as error bars. Age differences are most pronounced at Cz but are attenuated over more posterior sites. (B) Sagittal, coronal, and axial depiction of the association between gray matter volume and age (yellow), frontal (violet), and Cz (cyan) fast spindle density (separate voxel-vise multiple regression models for each predictor, controlled for total intracranial volume). In contrast to a global age-related decrease in gray matter volume (see Supplementary Figure 3, Supplementary Table 2), associations with spindle density are more specific. Whereas inter-individual differences in fast spindle density at Pz and Oz are not related to gray matter volume, fast spindle density at Cz shows a strong association with structural integrity in frontal and temporal brain regions (see Supplementary Figure 4, Supplementary Table 3). Clusters are displayed and considered significant at a FWE-corrected voxel threshold of *p* < .05, and a FWE-corrected cluster extent threshold of *p* < .05 and *k* >100. All clusters are corrected for nonstationary smoothness. See Supplementary Tables 3–5 for the cluster statistics and Supplementary Figures 4–6 for detailed depictions of the clusters. FWE: family-wise error; YA: younger adults; OA: older adults.

To derive a meaningful interpretation of the observed structure–function associations, one needs to consider that the extent to which we can detect oscillatory events on the scalp depends on both their generation and propagation. Thus, we can encounter the situation that oscillations are properly generated, but their propagation to certain brain areas is impaired by altered structural properties of the involved brain circuits (Landolt and Borbély, 2001; Martin et al., 2013; Nir et al., 2011; Sprecher et al., 2016). In line with this notion and the gray matter associations reported above, the maximum of fast spindle density over the posterior cortex has been interpreted in terms of aggravated fast spindle propagation to frontal and central brain regions that is severely affected by brain atrophy during aging. In contrast, the generation of sleep spindles is often regarded as less affected by aging (Crowley et al., 2002; Landolt and Borbély, 2001; Martin et al., 2013; Nir et al., 2011; Sprecher et al., 2016; see also Box 2). Still, we must acknowledge that the available cross-sectional correlative data neither allows us to disentangle whether differences in oscillatory components stem from disparities in their generation or their propagation, nor whether structural brain alterations cause, accompany, or result from alterations in sleep physiology. To solve these questions, longitudinal study designs that identify and track the sources of oscillatory components and link them to aging changes in structural brain integrity are required.

To conclude, the topographic distinctiveness of age-related sleep alterations is an important but often overlooked factor in age-comparative sleep studies. With the choice *where* to study age differences, age effects can either be maximized or reduced. Hence, deriving measures of EEG activity from only one derivation, as common in standard sleep recordings, should be avoided (Mander et al., 2017). To capture regional variation of age effects in the EEG, electrode setups that can capture hemispherical asymmetry (Sprecher et al., 2016) and allow for a high spatial resolution should be preferred (ideally at least a 32-electrode setup or even high density EEG). The fact that sleep parameters are selectively altered over specific brain regions, but not over other areas, casts doubt on the reported extent of age-related impairments in sleep processes. Brain atrophy likely results in a diminished and altered generation of certain sleep parameters (Dubé et al., 2015; Mander et al., 2013), but may also affect the propagation of electrical events that, per se, may be less prone to age-related changes. Future research should interpret the reported effects with care and provide solutions to disentangle age-related impairments in slow wave and spindle generation and propagation. Rather than placing a continued focus on the deficiency of sleep in old age, sleep studies require a shift towards maintained and intact sleep characteristics to determine altered or preserved functionality of sleep in old age.

## Conclusion

Fundamental changes in human sleep during aging are consistently reported and easy to observe – but, as discussed in this paper, it is nontrivial to quantify and measure them. Intact sleep and the functional succession of different sleep states is enabled by the balance of different neurotransmitter systems (Aston-Jones et al., 2007; Cirelli et al., 2005; Ramm, 1979; Steriade et al., 1990), and synergistic activity in different cortical and sub-cortical brain regions (Hobson and Pace-Schott, 2002; Pace-Schott and Hobson, 2002; Saletin et al., 2013; Steriade et al., 1990). Furthermore, it is embedded in a system of regulatory circadian and homeostatic processes (Achermann and Borbély, 1999; Dijk et al., 2000). Altogether, the diversity of regulatory mechanisms, reaching from genes and cells to neurotransmitters and brain networks (Hobson and Pace-Schott; Pace-Schott and Hobson, 2002), makes sleep susceptible to a wide range of possible interference. In consequence, the causes and the appearance of age-related changes in sleep are manifold and diverse (Dubé et al., 2015; Fogel et al., 2012, 2017; Mander et al., 2017; Muehlroth et al., 2019b; Zhong et al., 2019).

Standard criteria applied to measure these alterations are commonly optimized to capture inter-individual variance in healthy younger adults but can be severely biased when applied in other populations. As a result, not all age differences documented via the use of conventional analysis pipelines capture real alterations in underlying neural processing. We thus stress the need to develop individually adjusted analytic approaches in order to disentangle true age-dependent effects from age-independent inter-individual differences. But how could such an age-adjusted research process work?

We suggest that the basis for research on age-dependent differences in a given (neural) process is rooted in thorough theoretical considerations that incorporate the complex and dynamic nature of aging (e.g., Cabeza et al., 2018, 2005; Lindenberger et al., 2006) and the individual as the primary unit of analysis (Fisher et al., 2018; Grandy et al., 2017; Molenaar and Campbell, 2009; Nesselroade et al., 2007; Nesselroade and Molenaar, 2016; Rose et al., 2013). Ideally, such a research process starts out with an exact definition of a neural or physiological phenomenon that we intend to investigate. For instances we could ask whether the *intensity* of sleep (in terms of sleep depth) is reduced in old age. In a next step, potential indicators that mirror the phenomenon of interest along with its crucial features have to be considered and the most appropriate indicator(s) need(s) to be selected. In our example, we could decide to use SWA and slow oscillation density as markers of low-frequency oscillatory activity during sleep that index the depth of sleep (e.g., Achermann and Borbély, 1999; Bersagliere and Achermann, 2010; Blake and Gerard, 1937). Next, we need to determine how to best measure the respective indicator. Here, the individual has to be at the center of the analysis. Inter-individual differences that are irrelevant for our research question and thus introduce ‘noise’ (e.g., differences in anatomical skull properties) should be filtered out by adjusting analytic detection criteria at the level of the individual person (Karch et al., 2015; Molenaar and Campbell, 2009; Nesselroade et al., 2007; Nesselroade and Molenaar, 2016; Rose et al., 2013). In this way, we can create the conditions that facilitate an accurate interpretation of remaining inter-individual differences (Nesselroade et al., 2007). To be as unbiased as possible when comparing heterogeneous subject groups and individuals, we thus believe that research on sleep and aging should strive to identify the best possible analysis strategy to study a certain phenomenon of interest – even if this means moving away from conventional analytic approaches.

Complementarily, we stress the inevitable need to shift research foci from cross-sectional towards longitudinal studies. The currently available evidence cannot inform about ‘real’ age changes in sleep (Hertzog and Nesselroade, 2003; Hofer and Sliwinski, 2001; Li and Schmiedek, 2002; Lindenberger and Pötter, 1998; Lindenberger et al., 2011; Overton, 2010; Raz and Lindenberger, 2011). To disentangle adaptive from maladaptive aging processes, and to identify the lead-lag relations between age-related alterations in brain structure and sleep, study designs that follow the same individual across multiple measurement occasions are required (Hofer and Sliwinski, 2001; Li and Schmiedek, 2002; Lindenberger, 2014; Raz and Lindenberger, 2011).

Aside from that, we want to emphasize that age-dependent alterations in sleep physiology should not necessarily be interpreted in terms of ‘shortage’ or ‘deficiency.’ The brain is an *adaptive* organ that changes its structure and function in reaction to mismatches between environmental demands and organismic supplies (Lövden et al., 2010; Park and Reuter-Lorenz, 2009). First, this means that not all observed age differences in sleep necessarily indicate a loss (Baltes, 1987; Baltes et al., 1998). Reduced SWA, for instance, might merely be the consequence of reduced sleep pressure and an altered need for sleep in old age (Dijk et al., 2000). Second, due to the adaptive nature of the human brain, persons with very different developmental trajectories and physiological conditions can display similar sleep patterns (Muehlroth et al., 2019a). By identifying the similarities among individuals, and specifically among younger and older adults (Nesselroade et al., 2007), we can determine whether and how sleep is preserved in old age. Moreover, we can determine whether maintained sleep physiology is also reflected on the level of behavioral outcomes (Cabeza et al., 2018; Muehlroth et al., 2019a; Nyberg et al., 2012; Nyberg and Pudas, 2019; Park and Reuter-Lorenz, 2009). Research that shifts its focus towards the successful retention of sleep characteristics represents the foundation to gain a mechanistic understanding of the consequences entailed in altered sleep physiology. This is the basis for identifying the diagnostic value of sleep changes (Ferrarelli et al., 2019; Grandy et al., 2013; Prinz, 1977).

Finally, we want to point out that understanding sleep neurophysiological dynamics in their entirety ultimately requires taking a holistic view on sleep as alternating states of NREM and REM sleep (Conte and Ficca, 2013; Prerau et al., 2017; Scullin and Gao, 2018). Beyond a continued focus on NREM sleep, theories and analysis methods should take the cyclic nature of NREM and REM sleep into account and consider their dynamic interaction (e.g., by investigating stage transition rates, probabilities, and their temporal pattern; cf. Kishi et al., 2011; Schlemmer et al., 2015; Yetton et al., 2018).

Recent attempts to establish sleep as a novel biomarker and treatment target for Alzheimer’s disease have instantly evoked a massive rise in research on sleep and aging (Ju et al., 2014; Mander et al., 2016; Noble and Spires-Jones, 2019; Vaou et al., 2018). Methodological considerations, though, call for a clarification of *what* we want to measure and *how* we can best assess the respective phenomena. Improved methodological approaches will clearly speed up the search for a mechanistic understanding of the association between sleep and cognition in aged individuals.

### Box 3. Methodological advice for age-comparative sleep studies

1. Closely inspect your sleep EEG data visually. Knowing what the data looks like will ease your understanding of what to consider and what to adjust in your analyses.
2. Analyses should focus on micro-structural rather than macro-structural sleep features.
3. Define the neural process of interest as well as possible. Avoid proxies wherever possible.
4. Consider that seemingly similar measures (e.g., frequency, amplitude, density, power of the very same spindle event) can reflect very different physiological and neuronal properties. Be precise on which features you extract and interpret them carefully.
5. NREM sleep forms a continuum. Acknowledge that neurophysiological processes that are captured by sleep spindles and slow oscillations are present during both stage 2 sleep and SWS.
6. Use individually adjusted algorithms (with regard to amplitude and frequency) for sleep oscillation detection.
7. Use electrode setups that enable topographical comparisons. Try to incorporate the mechanisms that are driving topographical heterogeneity in age effects into your theories and methods (e.g., by carrying out structural brain imaging or source localization techniques in addition).
8. Instead of focusing on deficits only, consider what is intact or maintained in old age. Do your very best to apply age-adjusted methodology and comparisons.

## Acknowledgements

The authors thank Ann-Kathrin Joechner for helpful comments on an earlier version of the manuscript and Julia Delius for editorial assistance. We are grateful to everyone who contributed to the collection of data utilized in this article.

## Supplementary Methods

### Overview of data

In the present article, we report on data from 29 healthy younger (range: 19–28 years, *M*_age_ = 23.55 years, *SD*_age_ = 2.58; 16 females) and 34 healthy older adults (range: 63–74 years, *M*_age_ = 68.85 years, *SD*_age_ = 3.11; 15 females) whose sleep was monitored in two consecutive nights at their homes using ambulatory polysomnography (PSG). Analyses involving structural magnet resonance imaging (MRI) data, were based on data from 29 younger adults (*M*_age_ = 23.66 years, *SD*_age_ = 2.6; 17 female) and 32 of the older adults (*M*_age_ = 68.78 years, *SD*_age_ = 2.94; 14 female). All participants were right-handed native German speakers with no reported history of psychiatric or neurological disease, or any use of psychoactive medication. Older participants were screened for cognitive deficits using the Mini-Mental State Examination (MMSE; *M* = 29.24, *SD* = 1.12, range: 26–30; Folstein et al., 1975). General subjective sleep quality was assessed using the Pittsburgh Sleep Quality Index (PSQI; Buysse et al., 1989).

All displayed data were collected as part of a research project investigating age differences in the encoding, consolidation, and retrieval of associative memories (for more details on the experimental procedure, see Muehlroth et al., 2019a,b; Fandakova et al., 2018; Sander et al., 2019; Sommer et al., 2019). Participants’ sleep was monitored in two successive nights before and after engaging in an extensive associative learning task. An adaptation night prior to the first experimental night familiarized participants with the PSG procedure. MRI data were collected after the second experimental night. The research protocol was approved by the Ethics Committee of the *Deutsche Gesellschaft für Psychologie* (DGPs) and conducted at the *Max Planck Institute for Human Development* in Berlin. Informed written consent was obtained from all participants before study participation.

### Sleep data acquisition

Sleep was recorded in two consecutive nights using an ambulatory PSG device (SOMNOscreen plus; SOMNOmedics, Germany). Eight scalp electrodes were attached according to the international 10–20 system for electrode positioning (Fz, F3, F4, C3, C4, Cz, Pz, Oz; Jasper, 1958; Oostenveld and Praamstra, 2001) along with two electrodes on the mastoids A1 and A2, which later served as the offline reference. All impedances were kept below 6 kΩ. Data were recorded using Cz as the online reference for all EEG derivations and AFz as ground. Additionally, a bilateral electrooculogram (EOG) was assessed. Two submental electromyogram channels (EMG) were positioned left and right, below the labial angle, and referenced against one chin electrode. Electrical activity of the heart was recorded using two electrocardiogram channels (ECG). EEG channels were recorded between 0.2–75 Hz using a 16-bit analog-to-digital converter and a sampling rate of 128 Hz, with a manufacturers high-passed filter at 0.2 Hz and a low-pass filter at 35.0 Hz.

### EEG pre-processing

Data was first preprocessed using *BrainVision Analyzer 2.1* (Brain Products, Germany). Here, all EEG channels were re-referenced against the average of A1 and A2. To visually score the sleep data according to the standard criteria suggested by the *American Academy for Sleep Medicine* (AASM; Iber et al., 2007), the data were filtered using their recommended settings (EEG: 0.3–35 Hz; EOG: 0.3–35 Hz; EMG: 10–100 Hz). A Notch (50 Hz) filter was applied to the EMG and EOG derivations to remove external electrical interference. Afterwards, sleep stages 1 and 2, SWS, REM sleep, awakenings, and body movements were visually scored in 30-second epochs using *SchlafAus 1.4* (Gais & Werner, Lübeck, Germany). Total sleep time (TST) was calculated as the time spent in stage 1, stage 2, SWS, and REM sleep. Wake after sleep onset (WASO) was defined as the time participants were awake between sleep onset and final morning awakening.

Further analyses of the sleep EEG data were conducted using *Matlab R2014b* (Mathworks Inc., Sherbom, MA), the open-source toolbox *Fieldtrip* (Oostenveld et al., 2011), and custom-made code. Bad EEG channels were visually rejected. For the remaining channels, artefact detection was implemented on 1-second segments. Segments that were visually identified as body movement and those that exhibited amplitude differences of more than 500 μV were excluded. To further exclude segments that strongly deviated from the observed overall amplitude distribution, mean amplitude differences for each segment were *z*-standardized within each channel. Segments with a *z*-score of more than 5 in any of the channels were excluded. Code and sample data to reproduce the whole EEG analysis pipeline will be available on the Open Science Framework (https://osf.io/zur2b/).

### Slow oscillation detection

Slow oscillation detection at frontal electrodes was implemented based on previously established algorithms (Mölle et al., 2002; Ngo et al., 2013; cf. Muehlroth et al., 2019b). For NREM epochs (i.e., stage 2 sleep, and SWS), EEG data was band-pass filtered between 0.2 and 4 Hz using a 6th-order Butterworth filter (in forward and backward directions to prevent phase distortions). The whole signal was then divided into negative and positive half-waves that were separated by zero-crossings of the filtered signal. The combination of a negative half wave with the succeeding positive half wave was considered a putative slow oscillation when its frequency was between 0.5 and 1 Hz. The amplitude of each potential slow oscillation was calculated as the distance between the slow oscillation trough and positive peak, defined as its maximal negative and positive potential, respectively. Selection of real slow oscillations was then performed in two ways:

1. Adaptive amplitude thresholds were defined separately for each participant: Putative slow oscillations exceeding a trough of 1.25 times the mean trough of all putative slow oscillations as well as an amplitude of 1.25 times the average amplitude of all potential slow oscillations, were considered real slow oscillations (Ngo et al., 2013).
2. As an alternative detection criterion, fixed amplitude thresholds were defined: Putative slow oscillations exceeding a trough of –40 µV as well as an amplitude of 75 μV, were selected (Carrier et al., 2011; Dubé et al., 2015). Only artefact-free slow oscillations were considered in further analyses.

### Spindle detection

Spindles were detected using an established automated algorithm (Klinzing et al., 2016, 2018; Mölle et al., 2011). For NREM epochs (stage 2 sleep, and SWS), EEG data were band-pass filtered using a 6th-order Butterworth filter (in forward and backward directions) between 9 and 12.5 Hz for the detection of slow spindles, and between 12.5 and 16 Hz for fast spindles, respectively (Cox et al., 2017; Ujma et al., 2015). The root-mean-square (RMS) representation of the signal was calculated using a sliding window of 200 ms at each sample point. Afterwards, the RMS signal was smoothed by applying a moving average of 200 ms. To increase sensitivity and specificity of our algorithm (Coppieters ’t Wallant et al., 2016), we accounted for individual differences in EEG amplitude by anchoring the spindle identification to individually determined amplitude thresholds. A potential spindle was tagged if the amplitude of the smoothed RMS signal exceeded its mean by 1.5 SD of the filtered signal for 0.5 to 3 seconds. Spindles with boundaries closer than 0.25 seconds were eventually merged in the following way: In each processing run, starting with the smallest boundary difference, two spindles were merged if the resulting spindle event remained within the time limit of 3 seconds. As soon as a spindle was combined with another one, it could not be merged with another spindle in the same run again. Only if all boundary differences were revised and no further merging of putative spindles was possible, the new resulting spindle events were processed in a next run. Starting again with the smallest boundary difference, two putative spindles could be merged if the merged event remained within the required time limit. These runs were repeated until no further merging was possible. Finally, only spindles that did not overlap with previously marked artefact segments were considered real spindles.

### Spectral analysis

To derive whole-night characteristics of spectral EEG power (cf. Figure 2), we applied multitaper spectral analyses at electrode Cz (Mitra and Pesaran, 1999; Prerau et al., 2017) with frequency limits of 0.2 and 30 Hz and a frequency resolution of 1 Hz. Raw data was segmented into 30-second epochs that were spaced at 5-second intervals. A total of 29 discrete prolate spheroidal sequence (DPSS) tapers was applied. Power spectral analyses of the sleep EEG data were conducted, to obtain finer-grained indicators of neural processes during deep NREM sleep. Fast Fourier Transformation (FFT) with frequency limits between 0.5 and 30 Hz was applied on 5-second intervals using a Hanning function with no overlap and averaged across all NREM sleep (i.e., stage 2 sleep, and SWS). Log-transformed power spectra were derived for all channels. Slow-wave activity (SWA), defined as spectral power in the delta frequency range (0.5–4.5 Hz), was calculated for NREM epochs and normalized at the total power of the whole spectrum to minimize confounding effects and frequency-unspecific differences in EEG amplitudes (e.g., Dannhauer et al., 2011; Frodl et al., 2001; Leissner et al., 1970).

### Statistical analysis

Statistical analyses were conducted using RStudio 1.0.53 (RStudio, Inc., Boston, MA). For each subject, sleep estimates were averaged between the two experimental nights. Due to non-normal data distributions, we calculated non-parametric Mann-Whitney *U* Tests for independent samples and reported median and quartile values of the variables o identify age differences in sleep architecture and the expressed sleep oscillations between younger and older adults. Effect sizes were quantified by calculating Cohen’s *d* along with its 95 % confidence interval. Mixed factorial ANOVAs with the between factor AGE GROUP and the within factor CHANNEL were calculated to investigate the topographical differences in age-related sleep alterations. Post-hoc testing, again, was conducted using non-parametric Mann-Whitney *U* Tests for independent samples as well as Wilcoxon signed-rank tests for matched pairs. Associations between variables were estimated using Spearman’s rank-order correlation coefficient. The agreement between different sleep variables was quantified by calculating the *intraclass correlation coefficient* (ICC; Koo and Li, 2016) based on a two-way mixed effects model and consistency (= ICC(3,k)). To derive the test-retest reliability of the extracted sleep variables between the two consecutive nights, ICC calculation on the non-averaged sleep data was based on a two-way mixed effects model and absolute agreement (= ICC(2,k)). Significance levels were set to α = .05 and tested two-sided. We controlled statistical significance for multiple comparisons by Bonferroni-correcting the α-value and dividing it by the number of performed comparisons.

### Structural MRI data acquisition and analyses

Whole-brain MRI data was acquired with a Siemens Magnetom 3T TimTrio machine. A high-resolution T1-weighted MPRAGE sequence (TR = 2500 ms, TE = 4.77 ms, FOV = 256 mm, voxel size = 1 *×* 1 *×* 1 mm^3^) was collected from each participant. To quantify gray matter volume voxel-wise, we applied voxel-based morphometry (VBM; Ashburner and Friston, 2000) using *Statistical Parametric Mapping Software* (SPM12, http://www.fil.ion.ucl.ac.uk/spm) and the *Computational Anatomy Toolbox* (CAT 12, http://www.neuro.uni-jena.de/cat). Images were normalized to Montreal Neurological Institute (MNI) space and segmented into gray matter, white matter, and cerebrospinal fluid. Data were then modulated, i.e., corrected by the volume changes due to spatial normalization. For this, each voxel value was multiplied with the Jacobian determinant derived from the spatial normalization step. Afterwards images were smoothed with a 8 mm full width at half maximum (FWHM) kernel. Total intracranial volume (TIV) was estimated by summing volume of gray matter, white matter, and cerebrospinal fluid.

To derive the association between gray matter volume, age, and specific sleep variables, voxel-vise multiple regression models were implemented. For each subject, sleep estimates were averaged between the two experimental nights. All regression models were controlled for inter-individual differences in TIV by entering TIV as a covariate into the model. For masking, an absolute threshold of .02 was used. A family-wise error (FWE) correction was applied to all statistical maps, and α as set to .05. Further, we deployed a FWE-corrected cluster extent threshold of *p* < .05 and *k* >100 as well as a correction for nonstationary smoothness. Finally, we used the *xjView toolbox* (http://www.alivelearn.net/xjview/) to plot the resulting statistical *t*-maps onto the Ch2bet MNI template and to derive cluster labels.

## Supplementary Material

**Supplementary Figure 1.**
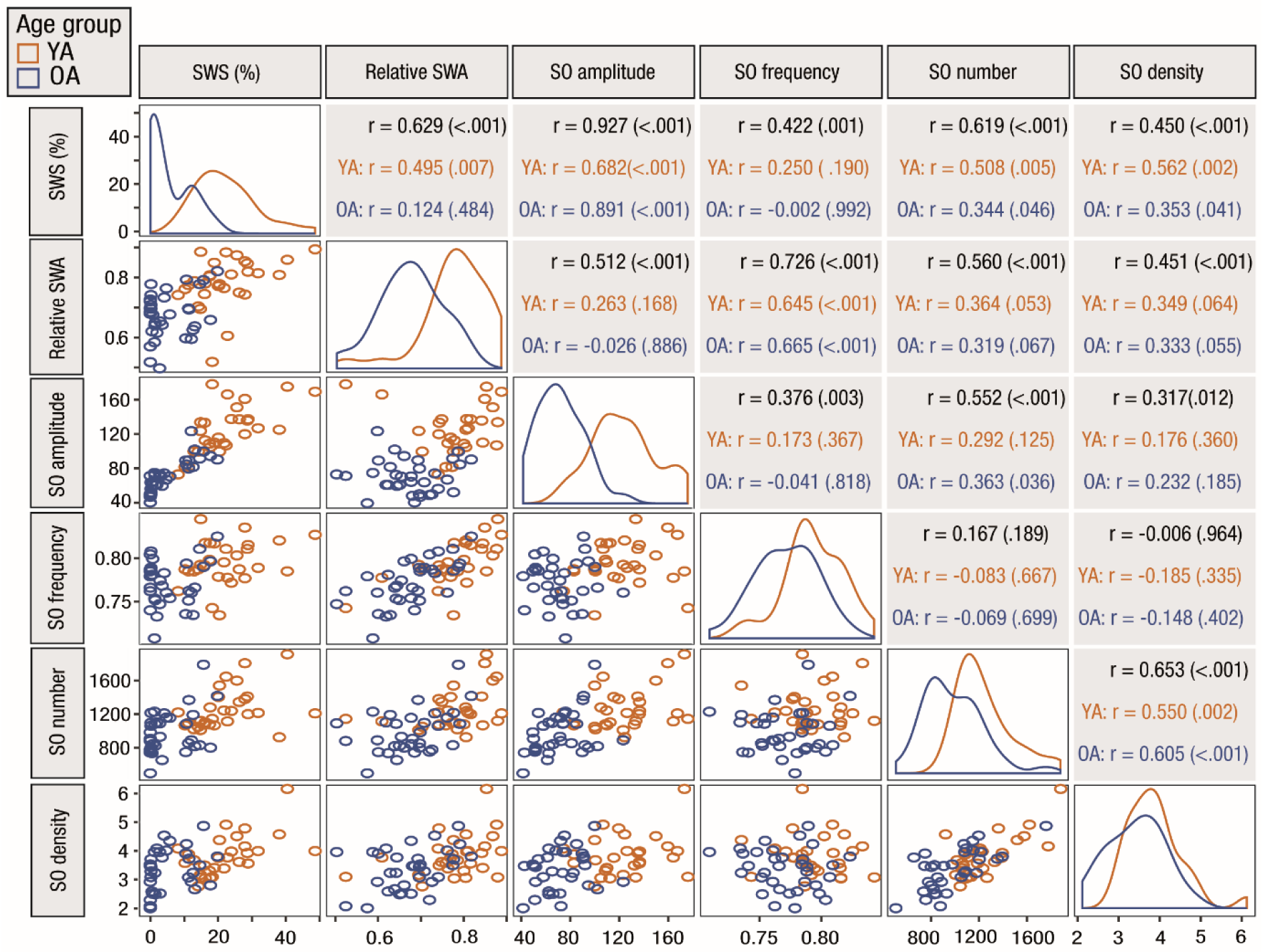
Correspondence between different indicators of slow oscillatory activity. Spearman’s rank-order correlation coefficients within and across age groups were calculated between the different indicators (corresponding *p*-values in brackets). All slow oscillation characteristics were derived using individually adjusted amplitude thresholds. The distribution of the respective variable within younger (orange) and older (blue) is displayed on the diagonal. Correlation coefficients and corresponding scatter plots are displayed in the upper right and lower left triangle respectively. OA: older adults; SO: slow oscillation; SWA: slow-wave activity; SWS: slow-wave sleep; YA: younger adults.

**Supplementary Figure 2.**
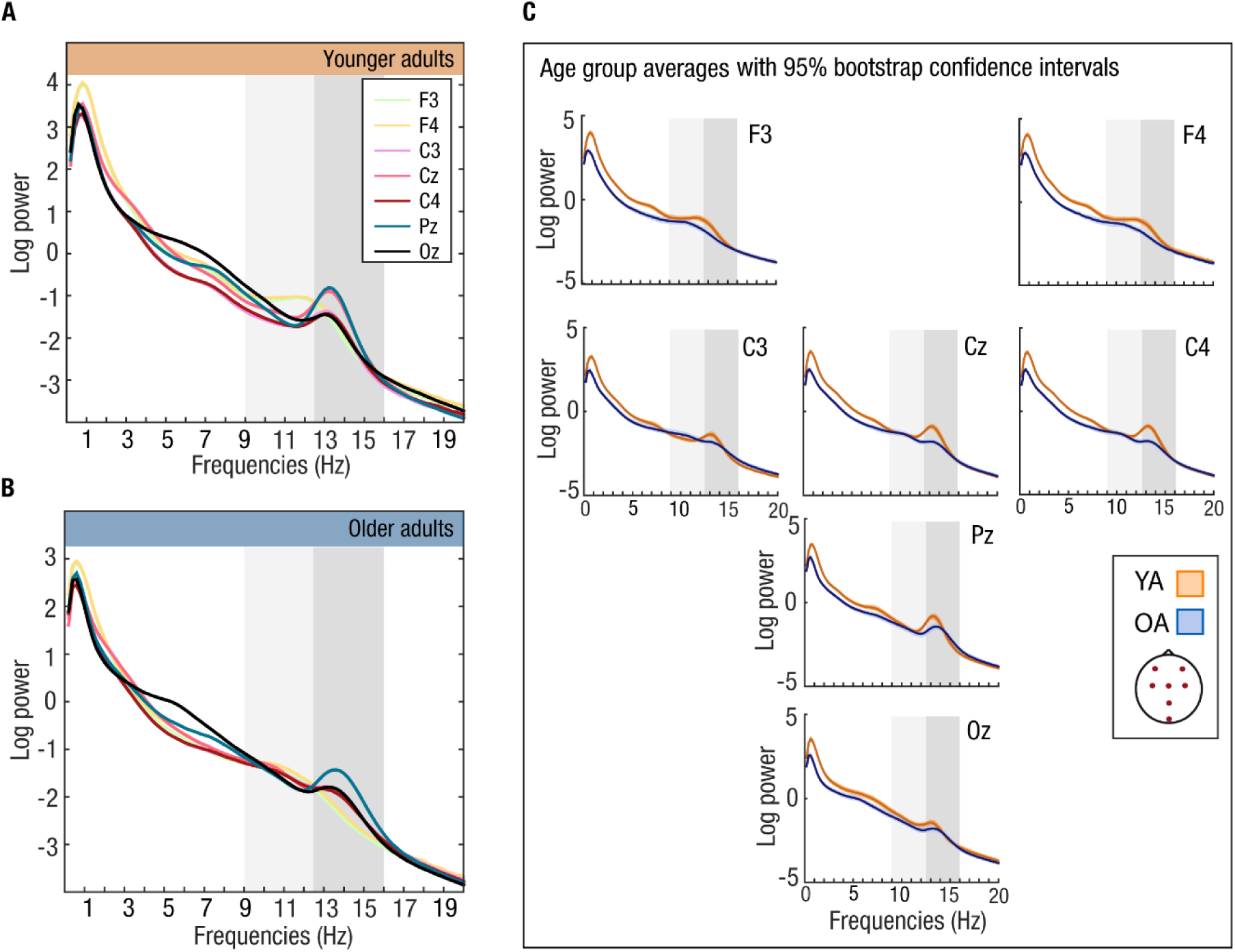
Log-transformed EEG power during NREM sleep. (A) In younger adults (YA) a clear peak in the fast spindle frequency range (12.5–16 Hz; shaded in dark gray) is observed in centroparietal electrodes. Frontal channels locally peak in lower frequencies at the transition from the slow (9–12.5 Hz, shaded in light gray) to fast spindle band. (B) In older adults (OA) we observe the same fast spindle power peak, but in Pz only. (C) For each age group the average power spectrum is depicted with its 95 % bootstrap confidence interval (500 bootstrap samples). Overall, the power spectra are flatter in OA (blue) than in YA (orange). YA show a stronger fast spindle power peak (shaded in dark grey) and higher power values in low frequencies (0–9 Hz) compared to OA. Note that, overall, confidence intervals are narrow. OA: older adults; YA: younger adults.

**Supplementary Figure 3.**
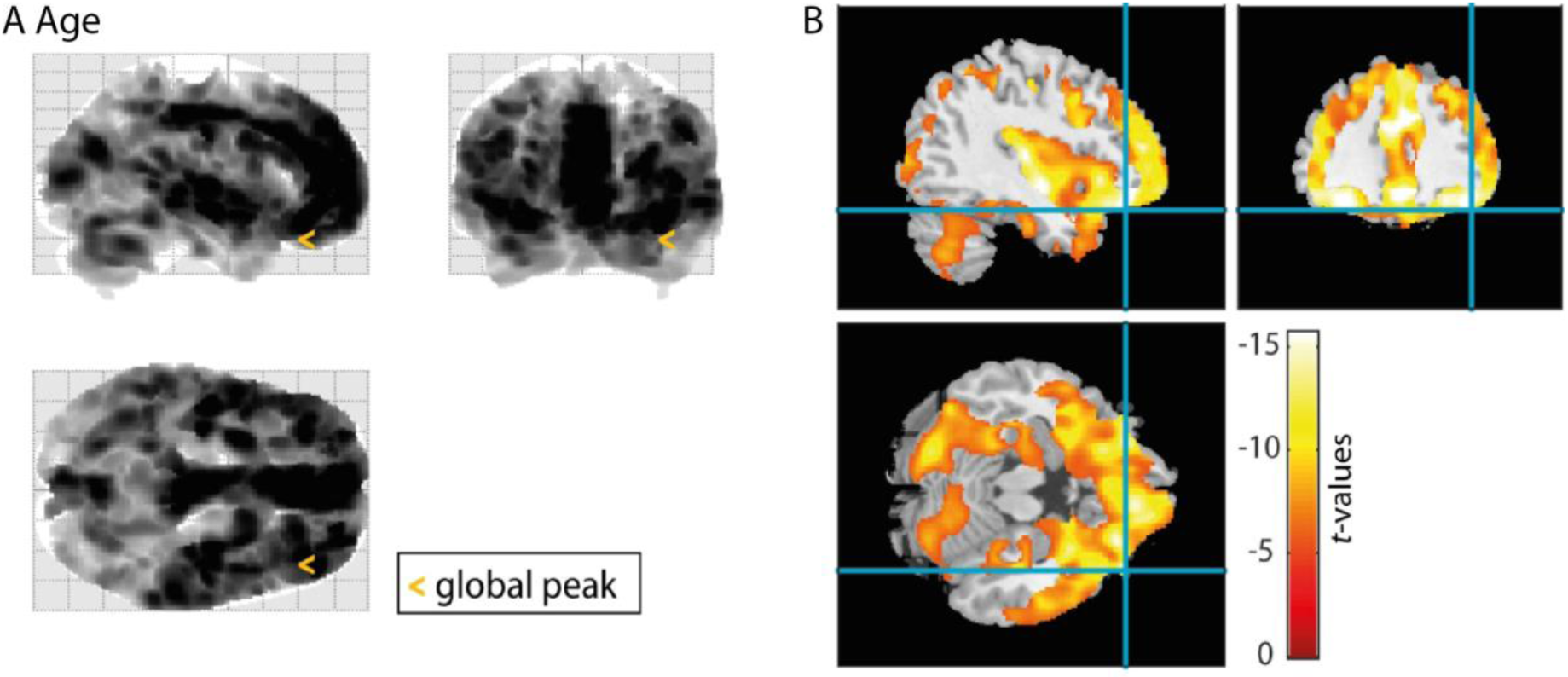
Significant negative cluster of the association between gray matter volume and age (multiple regression, controlled for total intracranial volume). The cluster’s peak intensity voxel is highlighted (glass brain (A): orange arrow; Ch2bet template brain (B): blue hair cross). The cluster is displayed and considered significant at a FWE-corrected voxel threshold of *p* < .05, and a FWE-corrected cluster extent threshold of *p* < .05 and *k* >100. It is corrected for nonstationary smoothness. See Supplementary Table 2 for the cluster statistics.

**Supplementary Figure 4.**
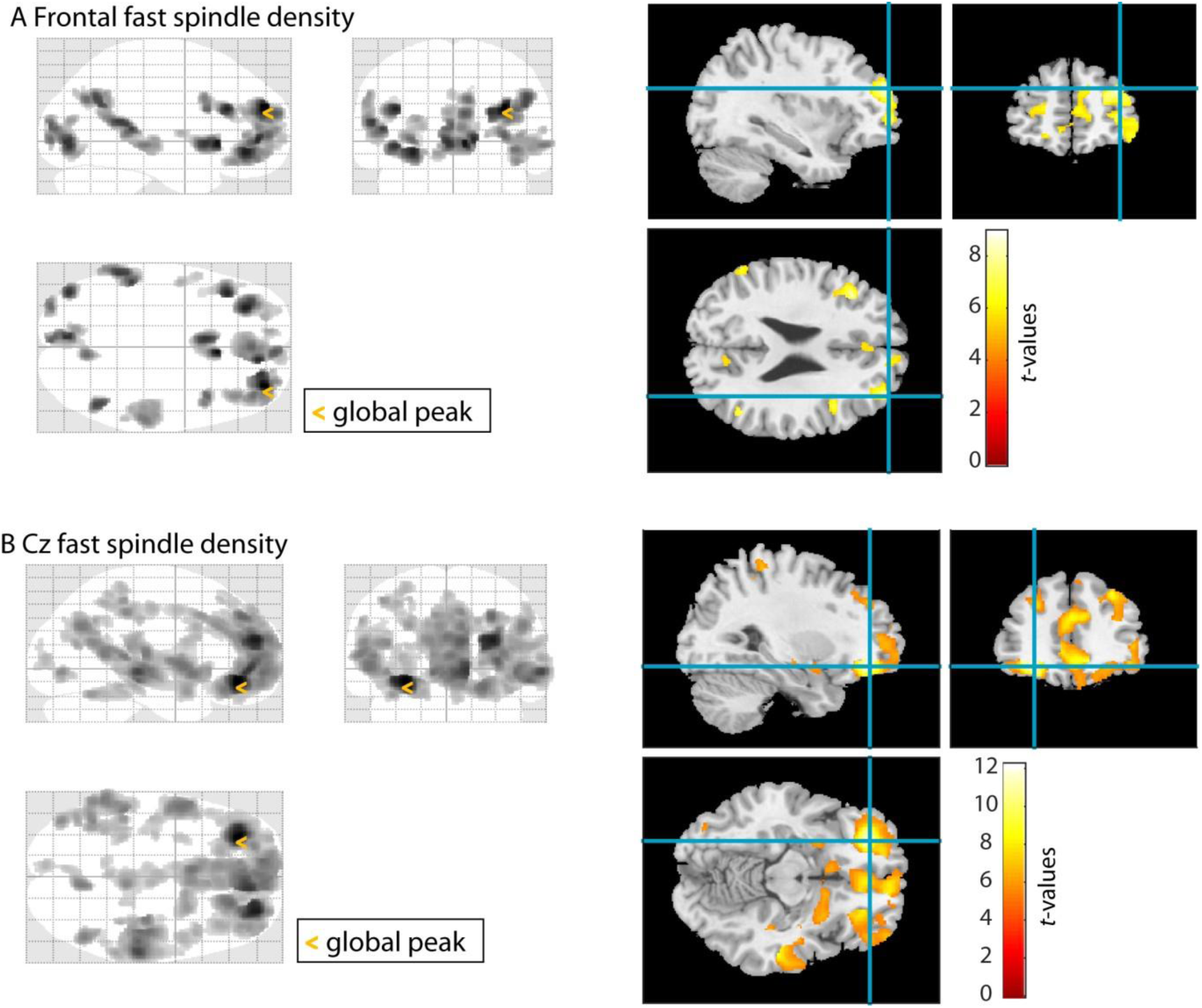
Significant negative clusters of the association between gray matter volume and fast spindle density at different electrode sites (multiple regression, controlled for total intracranial volume). The clusters’ peak intensity voxel is highlighted (glass brain: orange arrow; Ch2bet template brain: blue hair cross). Clusters are displayed and considered significant at a FWE-corrected voxel threshold of *p* < .05, and a FWE-corrected cluster extent threshold of *p* < .05 and *k* >100. All clusters are corrected for nonstationary smoothness. See Supplementary Table 3 for the cluster statistics.

**Supplementary Figure 5.**
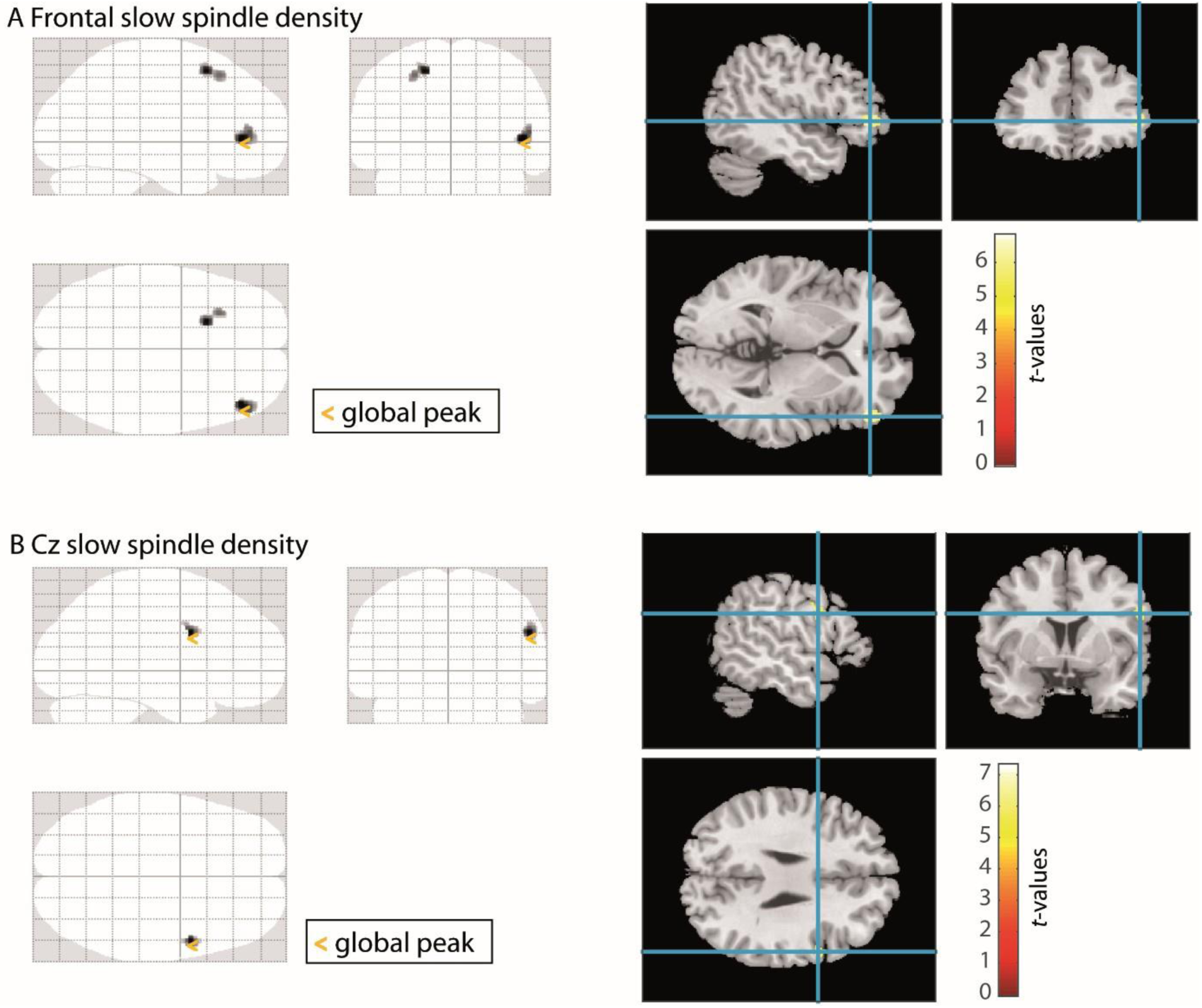
Significant negative clusters of the association between gray matter volume and slow spindle density at different electrode sites (multiple regression, controlled for total intracranial volume). The clusters’ global peak intensity voxel is highlighted (glass brain: orange arrow; Ch2bet template brain: blue hair cross). Clusters are displayed and considered significant at a FWE-corrected voxel threshold of *p* < .05, and a FWE-corrected cluster extent threshold of *p* < .05 and *k* >100. All clusters are corrected for nonstationary smoothness. See Supplementary Table 4 for the cluster statistics.

**Supplementary Figure 6.**
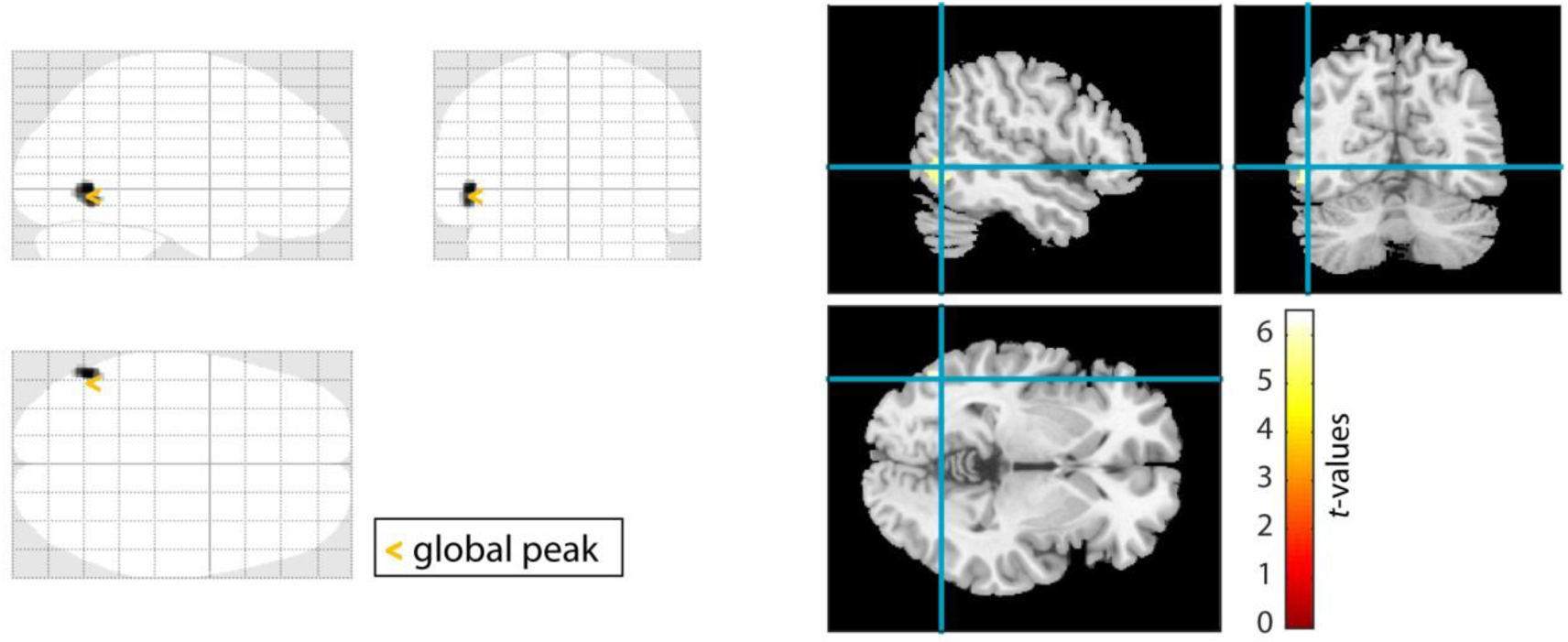
Significant negative cluster of the association between gray matter volume and slow oscillation density measured at Oz (multiple regression, controlled for total intracranial volume). The cluster’s global peak intensity voxel is highlighted (glass brain: orange arrow; Ch2bet template brain: blue hair cross). The cluster is displayed and considered significant at a FWE-corrected voxel threshold of *p* < .05, and a FWE-corrected cluster extent threshold of *p* < .05 and *k* >100. All clusters are corrected for nonstationary smoothness. See Supplementary Table 5 for the cluster statistics.

**Supplementary Figure 7.**
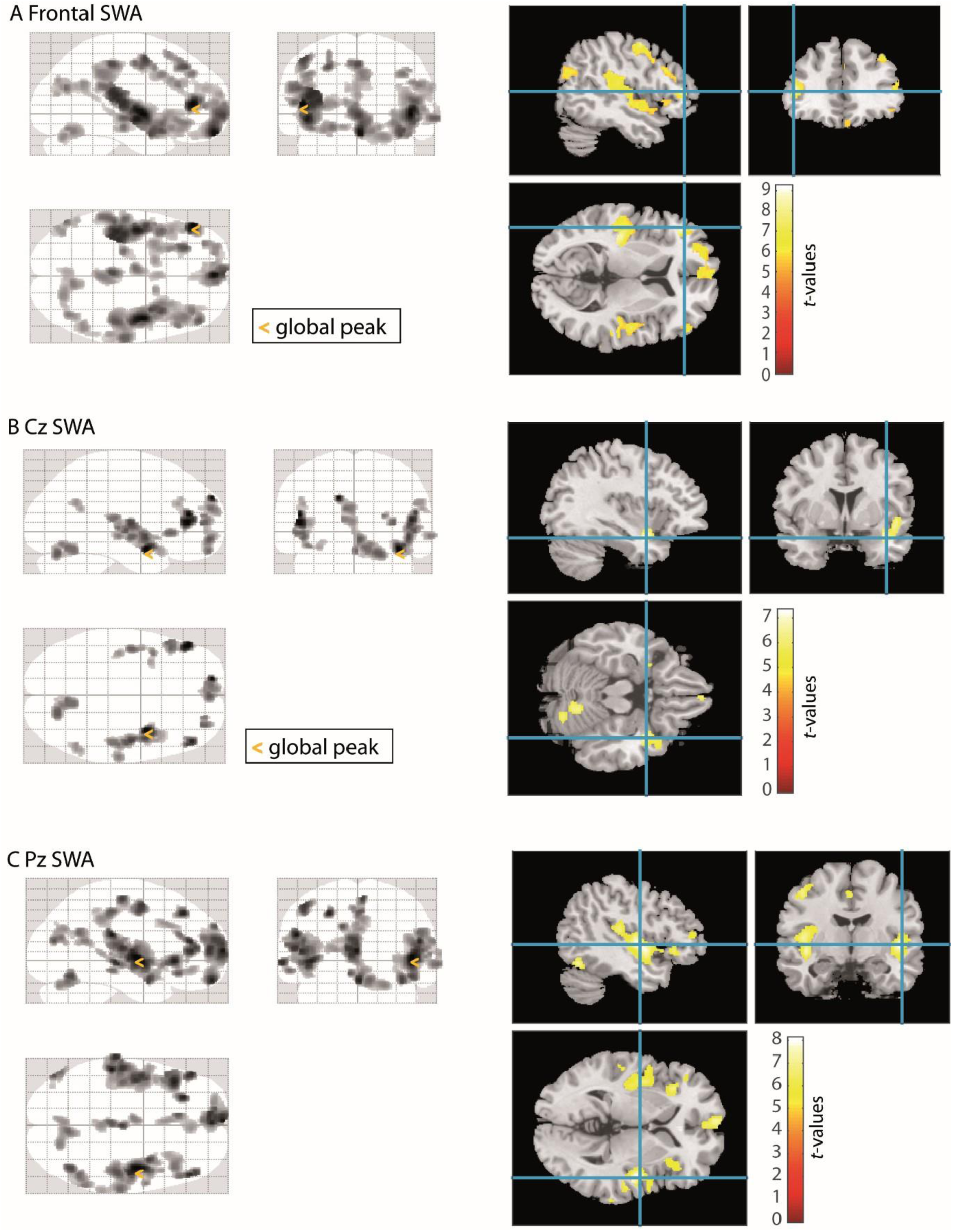
Significant negative clusters of the association between gray matter volume and SWA at different electrode sites (multiple regression, controlled for total intracranial volume). The clusters’ global peak intensity voxel is highlighted (glass brain: orange arrow; Ch2bet template brain: blue hair cross). Clusters are displayed and considered significant at a FWE-corrected voxel threshold of *p* < .05, and a FWE-corrected cluster extent threshold of *p* < .05 and *k* >100. All clusters are corrected for nonstationary smoothness. See Supplementary Table 6 for the cluster statistics.

**Supplementary Table 1.** Test-retest reliability for different markers of slow oscillatory activity

| | | ICC [95 % CI] | $F(df_1, df_2)$ | $p$ |
| --- | --- | --- | --- | --- |
| % SWS | Whole sample | .96 [.93; .98] | $F(62,62) = 25.17$ | < .001 |
| | YA | .9 [.78; .95] | $F(28,28) = 9.52$ | < .001 |
| | OA | .94 [.75; .98] | $F(33,33) = 23.83$ | < .001 |
| Relative SWA | Whole sample | .82 [.69; .90] | $F(62,62) = 6.19$ | < .001 |
| | YA | .83 [.64; .92] | $F(28,28) = 5.95$ | < .001 |
| | OA | .68 [.33; .85] | $F(33,33) = 3.69$ | < .001 |
| SO amplitude | Whole sample | .96 [.94; .98] | $F(62,62) = 25.96$ | < .001 |
| | YA | .94 [.87; .97] | $F(28,28) = 16.42$ | < .001 |
| | OA | .86 [.73; .93] | $F(33,33) = 7.35$ | < .001 |
| SO frequency | Whole sample | .89 [.82; .93] | $F(62,62) = 8.99$ | < .001 |
| | YA | .88 [.73; .94] | $F(28,28) = 7.98$ | < .001 |
| | OA | .85 [.74; .93] | $F(33,33) = 7.58$ | < .001 |
| SO number | Whole sample | .84 [.74; .91] | $F(62,62) = 6.35$ | < .001 |
| | YA | .83 [.64; .92] | $F(28,28) = 5.98$ | < .001 |
| | OA | .78 [.57; .89] | $F(33,33) = 4.61$ | < .001 |
| SO density | Whole sample | .88 [.79; .92] | $F(62,62) = 7.97$ | < .001 |
| | YA | .92 [.84; .96] | $F(28,28) = 13.08$ | < .001 |
| | OA | .81 [.62; .91] | $F(33,33) = 5.33$ | < .001 |

*Note.* Test-retest reliability was calculated between the two recorded experimental nights based on a two-way mixed effects model and absolute agreement. CI: confidence interval; ICC: intraclass correlation coefficient; OA: older adults; SO: slow oscillation; YA: younger adults.

**Supplementary Table 2.** Brain regions associated with age in a voxel-vise multiple regression, controlled for total intracranial volume.

| Anatomical peak region | Hemisphere | Peak intensity ($t$) | Peak MNI coordinates | | | Number of voxels |
| --- | --- | --- | --- | --- | --- | --- |
|  |  |  | x | y | z |  |
| Middle frontal gyrus, orbital part | R | -15.6536 | 39 | 36 | -19.5 | 167967 |

*Note.* The table lists cluster size along with the cluster's peak intensity (maximal $t$ -value, FWE-corrected) and corresponding MNI coordinates. Anatomical classification is based on the respective peak voxel. Reported clusters are significant with a FWE-corrected voxel threshold of $p < .05$ , a FWE-corrected cluster extent threshold of $p < .05$ and $k > 100$ and corrected for nonstationary smoothness. See Supplementary Figure 3 for the cluster depiction.

**Supplementary Table 3.**
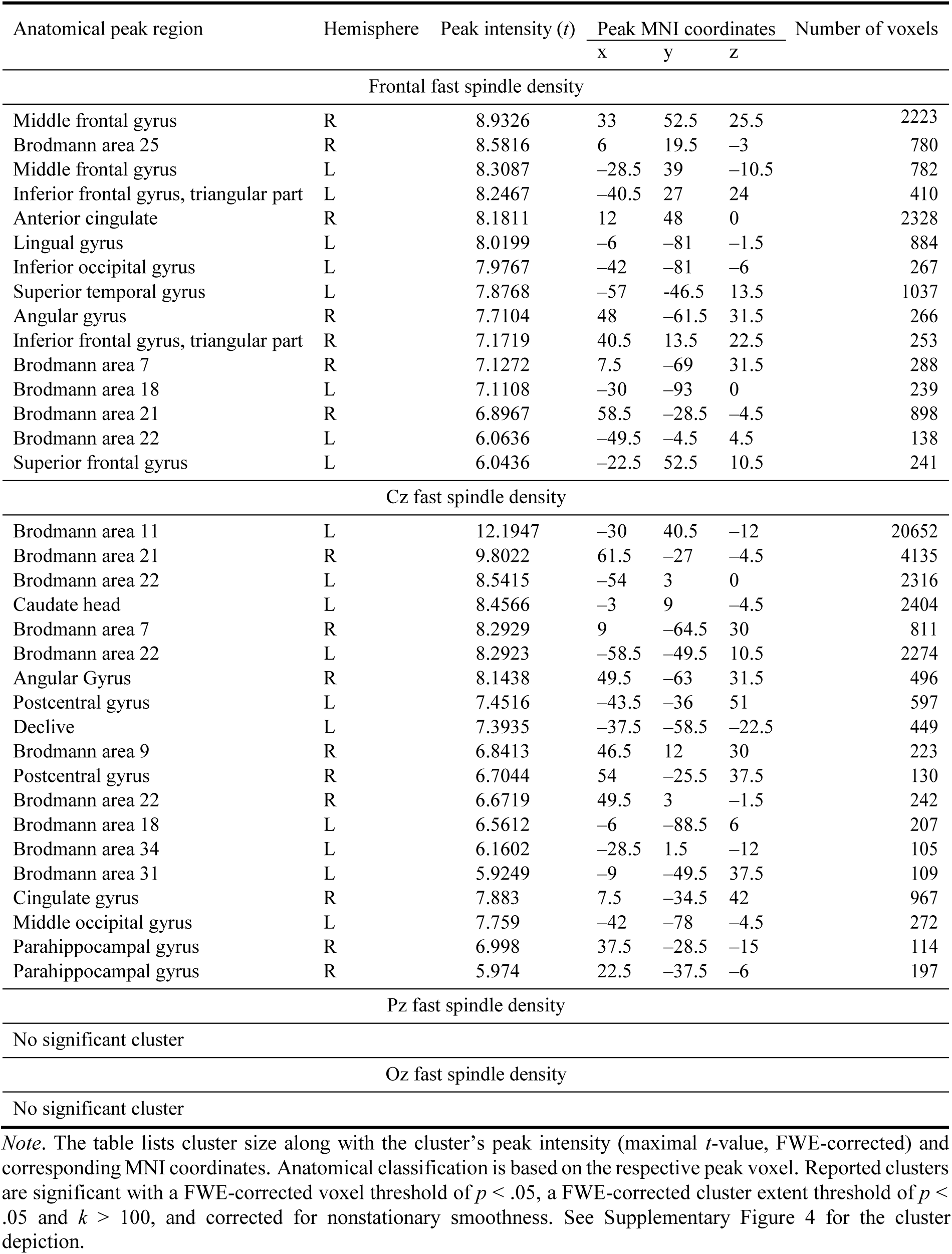
Brain regions associated with fast spindle density at different electrode sites (voxel-vise multiple regressions, controlled for total intracranial volume).

**Supplementary Table 4.** Brain regions associated with slow spindle density at different electrode sites (voxel-vise multiple regressions, controlled for total intracranial volume).

| Anatomical peak region | Hemisphere | Peak intensity (*t*) | Peak MNI coordinates |  |  | Number of voxels |
| --- | --- | --- | --- | --- | --- | --- |
|  |  |  | x | y | z |  |
| Frontal slow spindle density |  |  |  |  |  |  |
| Inferior frontal gyrus | R | 6.788 | 46.5 | 9 | 1.5 | 215 |
| Superior frontal gyrus | L | 6.7206 | −19.5 | 15 | 54 | 136 |
| Cz slow spindle density |  |  |  |  |  |  |
| Brodmann area 9 | R | 7.2931 | 52.5 | 4.5 | 28.5 | 100 |
| Pz slow spindle density |  |  |  |  |  |  |
| No significant cluster |  |  |  |  |  |  |
| Oz slow spindle density |  |  |  |  |  |  |
| No significant cluster |  |  |  |  |  |  |

*Note.* The table lists cluster size along with the cluster's peak intensity (maximal *t*-value, FWE-corrected) and corresponding MNI coordinates. Anatomical classification is based on the respective peak voxel. Reported clusters are significant with a FWE-corrected voxel threshold of $p < .05$ , a FWE-corrected cluster extent threshold of $p < .05$ and $k > 100$ , and corrected for nonstationary smoothness. See Supplementary Figure 5 for the cluster depiction.

**Supplementary Table 5.** Brain regions associated with slow oscillation density at different electrode sites (voxel-vise multiple regressions, controlled for total intracranial volume).

| Anatomical peak region | Hemisphere | Peak intensity (*t*) | Peak MNI coordinates |  |  | Number of voxels |
| --- | --- | --- | --- | --- | --- | --- |
|  |  |  | x | y | z |  |
| Frontal slow oscillation density |  |  |  |  |  |  |
| No significant cluster |  |  |  |  |  |  |
| Cz slow oscillation density |  |  |  |  |  |  |
| No significant cluster |  |  |  |  |  |  |
| Pz slow oscillation density |  |  |  |  |  |  |
| No significant cluster |  |  |  |  |  |  |
| Oz slow oscillation density |  |  |  |  |  |  |
| Middle temporal gyrus | L | 6.4661 | −51 | −63 | −1.5 | 155 |

*Note.* The table lists cluster size along with the cluster's peak intensity (maximal *t*-value, FWE-corrected) and corresponding MNI coordinates. Anatomical classification is based on the respective peak voxel. Reported clusters are significant with a FWE-corrected voxel threshold of $p < .05$ , a FWE-corrected cluster extent threshold of $p < .05$ and $k > 100$ , and corrected for nonstationary smoothness. See Supplementary Figure 6 for the cluster depiction.

**Supplementary Table 6.**
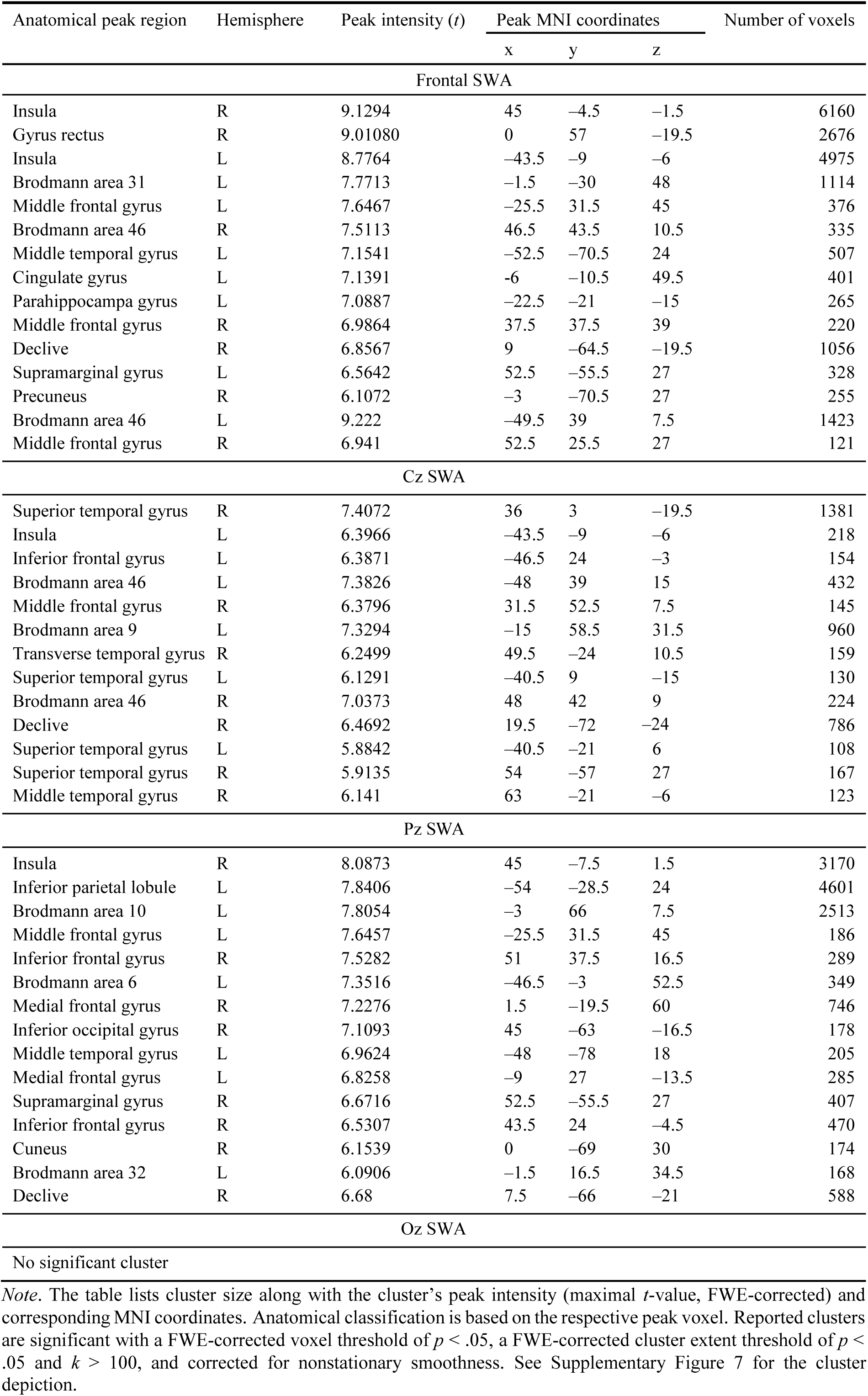
Brain regions associated with SWA at different electrode sites (voxel-vise multiple regressions, controlled for total intracranial volume).

| Anatomical peak region | Hemisphere | Peak intensity (*t*) | Peak MNI coordinates |  |  | Number of voxels |
| --- | --- | --- | --- | --- | --- | --- |
|  |  |  | x | y | z |  |
| Frontal SWA |  |  |  |  |  |  |
| Insula | R | 9.1294 | 45 | −4.5 | −1.5 | 6160 |
| Gyrus rectus | R | 9.01080 | 0 | 57 | −19.5 | 2676 |
| Insula | L | 8.7764 | −43.5 | −9 | −6 | 4975 |
| Brodmann area 31 | L | 7.7713 | −1.5 | −30 | 48 | 1114 |
| Middle frontal gyrus | L | 7.6467 | −25.5 | 31.5 | 45 | 376 |
| Brodmann area 46 | R | 7.5113 | 46.5 | 43.5 | 10.5 | 335 |
| Middle temporal gyrus | L | 7.1541 | −52.5 | −70.5 | 24 | 507 |
| Cingulate gyrus | L | 7.1391 | −6 | −10.5 | 49.5 | 401 |
| Parahippocampa gyrus | L | 7.0887 | −22.5 | −21 | −15 | 265 |
| Middle frontal gyrus | R | 6.9864 | 37.5 | 37.5 | 39 | 220 |
| Declive | R | 6.8567 | 9 | −64.5 | −19.5 | 1056 |
| Supramarginal gyrus | L | 6.5642 | 52.5 | −55.5 | 27 | 328 |
| Precuneus | R | 6.1072 | −3 | −70.5 | 27 | 255 |
| Brodmann area 46 | L | 9.222 | −49.5 | 39 | 7.5 | 1423 |
| Middle frontal gyrus | R | 6.941 | 52.5 | 25.5 | 27 | 121 |
| Cz SWA |  |  |  |  |  |  |
| Superior temporal gyrus | R | 7.4072 | 36 | 3 | −19.5 | 1381 |
| Insula | L | 6.3966 | −43.5 | −9 | −6 | 218 |
| Inferior frontal gyrus | L | 6.3871 | −46.5 | 24 | −3 | 154 |
| Brodmann area 46 | L | 7.3826 | −48 | 39 | 15 | 432 |
| Middle frontal gyrus | R | 6.3796 | 31.5 | 52.5 | 7.5 | 145 |
| Brodmann area 9 | L | 7.3294 | −15 | 58.5 | 31.5 | 960 |
| Transverse temporal gyrus | R | 6.2499 | 49.5 | −24 | 10.5 | 159 |
| Superior temporal gyrus | L | 6.1291 | −40.5 | 9 | −15 | 130 |
| Brodmann area 46 | R | 7.0373 | 48 | 42 | 9 | 224 |
| Declive | R | 6.4692 | 19.5 | −72 | −24 | 786 |
| Superior temporal gyrus | L | 5.8842 | −40.5 | −21 | 6 | 108 |
| Superior temporal gyrus | R | 5.9135 | 54 | −57 | 27 | 167 |
| Middle temporal gyrus | R | 6.141 | 63 | −21 | −6 | 123 |
| Pz SWA |  |  |  |  |  |  |
| Insula | R | 8.0873 | 45 | −7.5 | 1.5 | 3170 |
| Inferior parietal lobule | L | 7.8406 | −54 | −28.5 | 24 | 4601 |
| Brodmann area 10 | L | 7.8054 | −3 | 66 | 7.5 | 2513 |
| Middle frontal gyrus | L | 7.6457 | −25.5 | 31.5 | 45 | 186 |
| Inferior frontal gyrus | R | 7.5282 | 51 | 37.5 | 16.5 | 289 |
| Brodmann area 6 | L | 7.3516 | −46.5 | −3 | 52.5 | 349 |
| Medial frontal gyrus | R | 7.2276 | 1.5 | −19.5 | 60 | 746 |
| Inferior occipital gyrus | R | 7.1093 | 45 | −63 | −16.5 | 178 |
| Middle temporal gyrus | L | 6.9624 | −48 | −78 | 18 | 205 |
| Medial frontal gyrus | L | 6.8258 | −9 | 27 | −13.5 | 285 |
| Supramarginal gyrus | R | 6.6716 | 52.5 | −55.5 | 27 | 407 |
| Inferior frontal gyrus | R | 6.5307 | 43.5 | 24 | −4.5 | 470 |
| Cuneus | R | 6.1539 | 0 | −69 | 30 | 174 |
| Brodmann area 32 | L | 6.0906 | −1.5 | 16.5 | 34.5 | 168 |
| Declive | R | 6.68 | 7.5 | −66 | −21 | 588 |
| Oz SWA |  |  |  |  |  |  |
| No significant cluster |  |  |  |  |  |  |

*Note.* The table lists cluster size along with the cluster's peak intensity (maximal *t*-value, FWE-corrected) and corresponding MNI coordinates. Anatomical classification is based on the respective peak voxel. Reported clusters are significant with a FWE-corrected voxel threshold of $p < .05$ , a FWE-corrected cluster extent threshold of $p < .05$ and $k > 100$ , and corrected for nonstationary smoothness. See Supplementary Figure 7 for the cluster depiction.

